# Semantic reconstruction of continuous language from non-invasive brain recordings

**DOI:** 10.1101/2022.09.29.509744

**Authors:** Jerry Tang, Amanda LeBel, Shailee Jain, Alexander G. Huth

## Abstract

A brain-computer interface that decodes continuous language from non-invasive recordings would have many scientific and practical applications. Currently, however, decoders that reconstruct continuous language use invasive recordings from surgically implanted electrodes^1–3^, while decoders that use non-invasive recordings can only identify stimuli from among a small set of letters, words, or phrases^4–7^. Here we introduce a non-invasive decoder that reconstructs continuous natural language from cortical representations of semantic meaning^8^ recorded using functional magnetic resonance imaging (fMRI). Given novel brain recordings, this decoder generates intelligible word sequences that recover the meaning of perceived speech, imagined speech, and even silent videos, demonstrating that a single language decoder can be applied to a range of semantic tasks. To study how language is represented across the brain, we tested the decoder on different cortical networks, and found that natural language can be separately decoded from multiple cortical networks in each hemisphere. As brain-computer interfaces should respect mental privacy^9^, we tested whether successful decoding requires subject cooperation, and found that subject cooperation is required both to train and to apply the decoder. Our study demonstrates that continuous language can be decoded from non-invasive brain recordings, enabling future multipurpose brain-computer interfaces.

## Main Text

Previous brain-computer interfaces have demonstrated that speech articulation^1^ and other motor signals can be decoded from intracranial recordings to restore communication to people who have lost the ability to speak^2, 3^. While effective, these decoders require invasive neurosurgery, making them unsuitable for most other uses. Non-invasive language decoders could be more widely adopted, and may eventually help all people interact with technological devices through thought^4–7^.

We introduce a decoder that takes non-invasive fMRI brain recordings and reconstructs arbitrary stimuli that the subject is hearing or imagining in continuous natural language. To accomplish this we needed to overcome one major obstacle: the low temporal resolution of fMRI. While fMRI has excellent spatial specificity, the blood-oxygen-level-dependent (BOLD) signal that it measures is notoriously slow—an impulse of neural activity causes BOLD to rise and fall over approximately 10 seconds^10^. For naturally spoken English (over 2 words per second), this means that each brain image can be affected by over 20 words. Decoding continuous language thus requires solving an ill-posed inverse problem, as there are many more words to decode than brain images. Our decoder accomplishes this by guessing candidate word sequences, scoring the likelihood that each candidate evoked the recorded brain responses, and then selecting the best candidate.

To compare word sequences to a subject’s brain responses, we trained an encoding model^8^ that predicts how that subject’s brain responds to phrases in natural language. This model can score the likelihood that the subject is hearing or imagining a candidate sequence by measuring how well the recorded brain responses match the predicted brain responses^11, 12^. To learn how the brain responds to a wide range of phrases, we recorded responses while the subject listened to sixteen hours of naturally spoken narrative stories, yielding over five times more data than the typical language fMRI experiment. We trained an encoding model on this dataset by extracting semantic features that capture the meaning of stimulus phrases^13–17^, and then using linear regression to model how the semantic features influence brain responses (Fig. 1a). Given any word sequence, this encoding model predicts how the subject’s brain would respond.

**Fig. 1.**
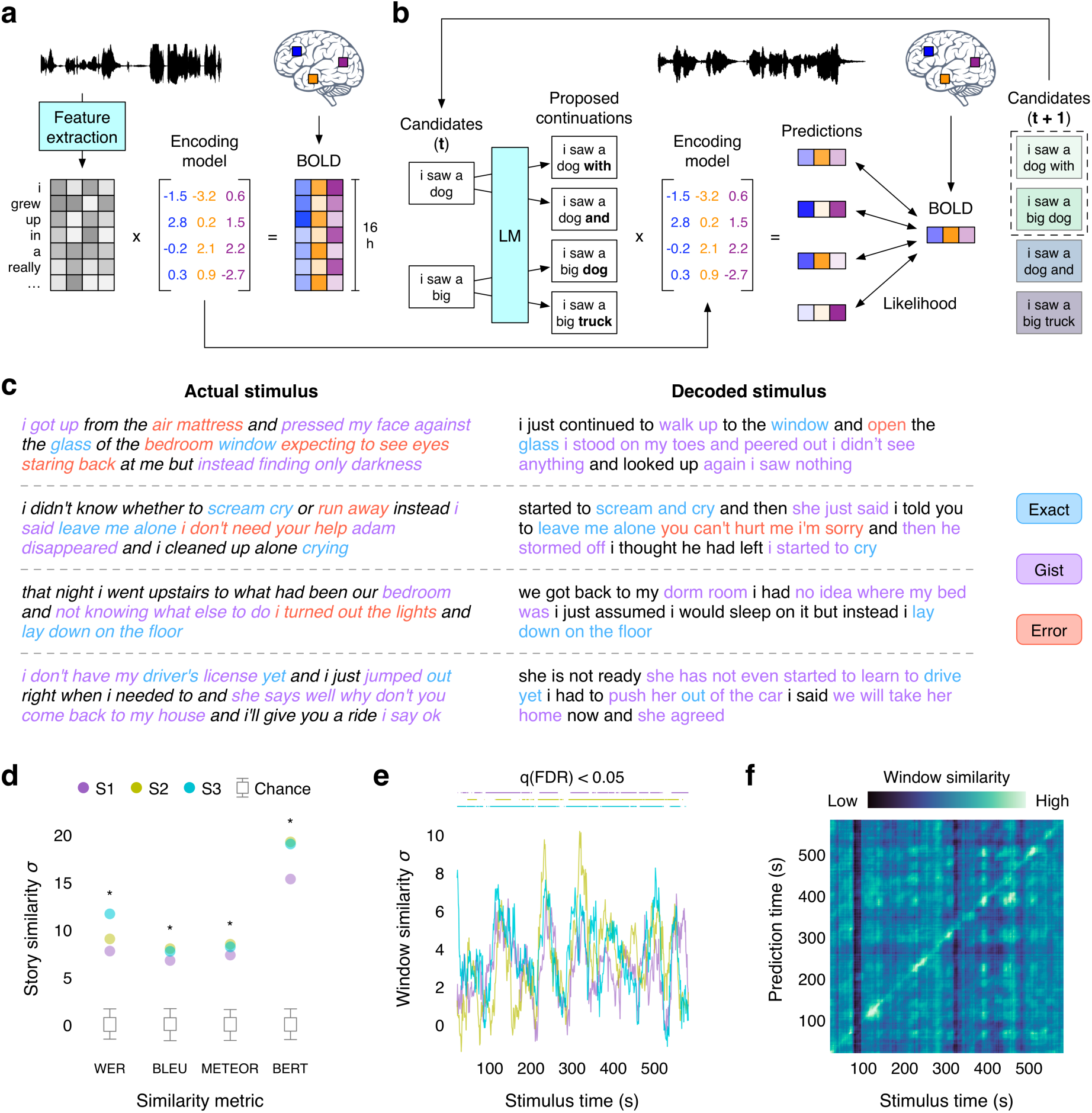
Language decoder. (**a**) BOLD fMRI responses were recorded while three subjects listened to 16 h of narrative stories. An encoding model was estimated for each subject to predict brain responses from semantic features of stimulus words. (**b**) To reconstruct language from novel brain recordings, the decoder maintains a set of candidate word sequences. When new words are detected, a language model (LM) proposes continuations for each sequence and the encoding model scores the likelihood of the recorded brain responses under each continuation. The most likely continuations are retained. (**c**) Decoders were evaluated on single-trial brain responses recorded while subjects listened to test stories that were not used for model training. Segments from four test stories are shown alongside decoder predictions for one subject. The decoder exactly reproduces some words and phrases, and captures the gist of many more. (**d**) Decoder predictions for a test story were significantly more similar to the actual stimulus words than expected by chance under a range of language similarity metrics (* indicates *q*(FDR) < 0.05 for all subjects, one-sided nonparametric test). To compare across metrics, results are shown as standard deviations away from the mean of the null distribution (see Methods). Boxes indicate the interquartile range of the null distribution; whiskers indicate the 5th and 95th percentiles. (**e**) For most time points, decoding scores were significantly higher than expected by chance (*q*(FDR) < 0.05, one-sided nonparametric test) under the BERTScore metric. (**f**) Identification accuracy for one subject. The brightness at (𝑖, 𝑗) reflects the similarity between the 𝑖th second of the prediction and the 𝑗th second of the actual stimulus. Identification accuracy was significantly higher than expected by chance (*p* < 0.05, one-sided permutation test).

In theory, we could identify the most likely stimulus that the subject is hearing or imagining by comparing the recorded brain responses to encoding model predictions for every possible word sequence^11, 12^. However, the number of possible word sequences is far too large for this approach to be practical, and the vast majority of those sequences do not resemble natural language. To restrict the candidate sequences to well-formed English, we used a generative neural network language model^18^ that was trained on a large dataset of natural English word sequences. Given any word sequence, this language model predicts the words that could come next.

Yet even with the constraints imposed by the language model, it is computationally infeasible to generate and score all candidate sequences. To efficiently search the space of word sequences, we used a beam search algorithm^19^ that generates candidate sequences word by word. In beam search, the decoder maintains a beam containing the 𝑘 most likely candidate sequences at any given time. When new words are detected based on brain activity in auditory and speech regions (see Methods), the language model generates continuations for each candidate sequence in the beam. The encoding model then scores the likelihood that each continuation evoked the recorded brain responses, and the 𝑘 most likely continuations are retained in the beam for the next timestep (Fig. 1b). This process continually approximates the stimulus that the subject is hearing or imagining across an arbitrary amount of time.

We trained decoders for three subjects and evaluated each subject’s decoder on separate, single-trial brain responses that were recorded while that subject listened to novel test stories that were not used for model training. Since our decoder represents language using semantic features rather than motor or acoustic features, the decoder predictions should capture the meaning of the stimuli. Results show that the decoded word sequences captured not only the meaning of the stimuli, but often even recovered exact words and phrases (Fig. 1c; Supplementary Table 1). To quantify decoding performance, we compared decoded and actual word sequences for one test story (1,800 words) using several language similarity metrics (see Methods). Standard metrics like word error rate (WER), BLEU, and METEOR measure the number of words shared by two sequences. However, because different words can convey the same meaning—for instance “we were busy” and “we had a lot of work”—we also used BERTScore, a newer method which uses machine learning to quantify whether two sequences share a meaning. Story decoding performance was significantly higher than expected by chance under each metric but particularly BERTScore (*q*(FDR) < 0.05, one-sided nonparametric test; Fig. 1d). Most time-points in the story (72-82%) had a significantly higher BERTScore than expected by chance (Fig. 1e) and could be identified from other time-points (mean percentile rank = 0.85-0.90) based on BERTScore similarities between the decoded and actual words (Fig. 1f; Extended Data Fig. 1a).

## Decoding across cortical networks

The decoding results shown in Figure 1 used responses from many different cortical regions to achieve good performance. This builds upon earlier reports that most parts of cortex contain some representations of language meaning^8, 20^. However, it is unclear which cortical networks represent language in sufficient detail to decode complete word sequences^6^, and whether different networks—or hemispheres—play complementary or redundant roles in language processing^21, 22^. To answer these questions, we partitioned brain data into three cortical networks—the classical language network^23^, the parietal-temporal-occipital association network^20^, and the prefrontal network—and separately decoded from each network in each hemisphere (Fig. 2a; Extended Data Fig. 1b).

**Fig. 2.**
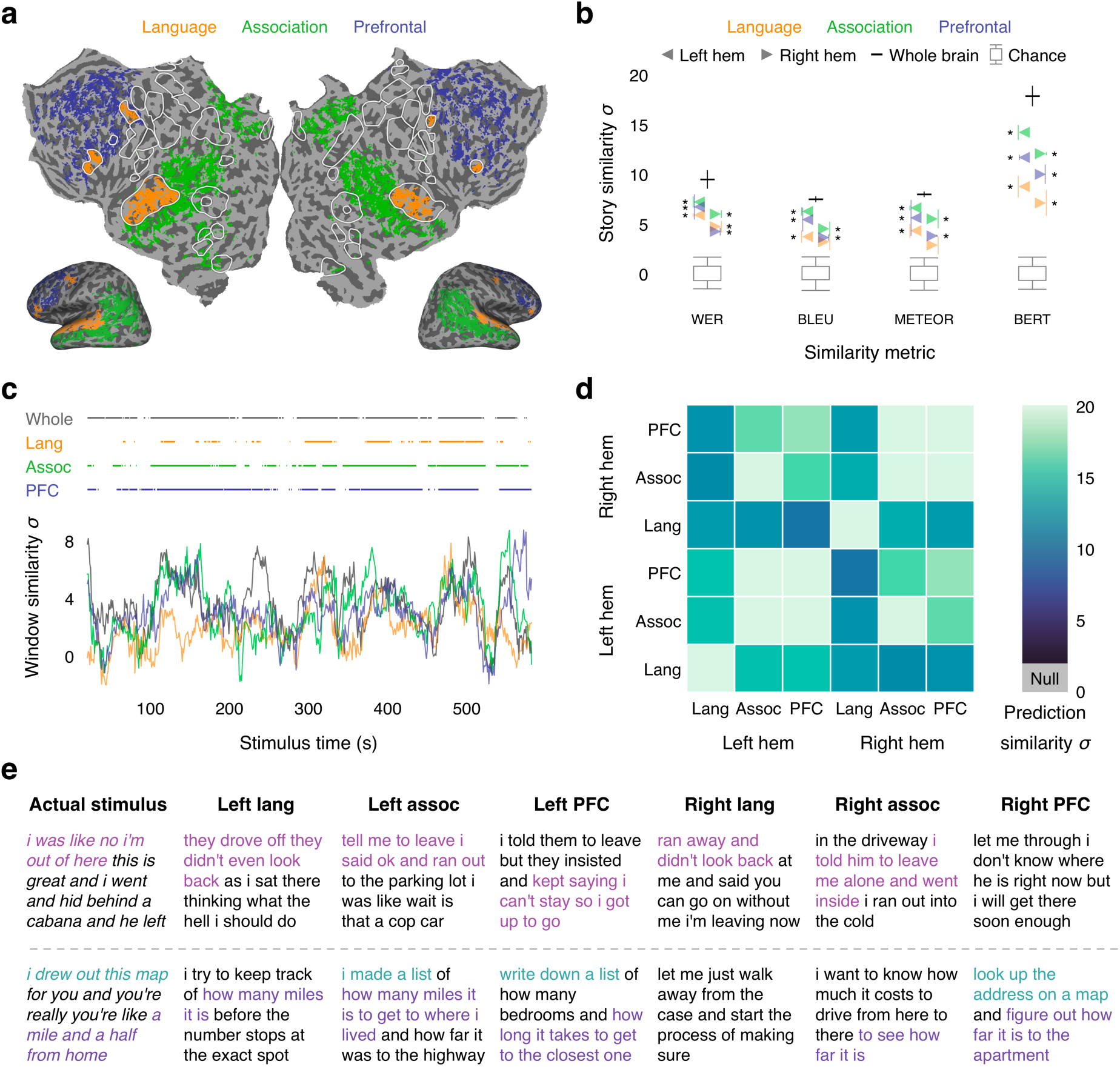
Decoding across cortical networks. (**a**) Cortical networks for one subject. Brain data used for decoding (colored regions) were partitioned into the classical language network, the parietal-temporal-occipital association network, and the prefrontal network (PFC). (**b**) Decoder predictions from each network in each hemisphere were significantly more similar to the actual stimulus words than expected by chance under most metrics (* indicates *q*(FDR) < 0.05 for all subjects, one-sided nonparametric test). Error bars indicate standard error of the mean (*n* = 3 subjects). Boxes indicate the interquartile range of the null distribution; whiskers indicate the 5th and 95th percentiles. (**c**) Decoding performance time-course from each network for one subject. Horizontal lines indicate when decoding performance was significantly higher than expected by chance under the BERTScore metric (*q*(FDR) < 0.05, one-sided nonparametric test). Most time-points that were significantly decoded from the whole brain were also significantly decoded from association and prefrontal networks. (**d**) Decoder predictions were compared across networks and hemispheres. Each pair of predictions were more similar than expected by chance (*q*(FDR) < 0.05, two-sided nonparametric test). (**e**) Segments from a test story are shown alongside decoder predictions from each network in each hemisphere for one subject. Colors indicate corresponding phrases. These results demonstrate that cortical networks contain redundant representations of natural language.

Decoder predictions from each network in each hemisphere were significantly more similar to the actual stimulus than expected by chance (*q*(FDR) < 0.05, one-sided nonparametric test; Fig. 2b). Notably, we successfully decoded continuous language from association and prefrontal networks without using any information from the classical language network except to detect when new words occurred. These results underscore the role of bilateral domain-general semantic regions in representing natural language^8, 24^.

One possible explanation for the successful decoding from multiple networks is that these networks encode complementary representations—such as different semantic categories—in a modular organization. If this were the case, different parts of the stimulus may be decodable from each individual network, but the full stimulus should only be decodable from the whole brain. Alternatively, the different networks might encode redundant representations. If this were the case, the full stimulus may be separately decodable from each individual network.

To differentiate these possibilities, we first computed the time-course of decoding performance for each network, and found that most time-points that were significantly decoded from the whole brain could also be decoded from association (77-83%) and prefrontal (54-82%) networks (Fig. 2c; Extended Data Fig. 1c). We then compared decoder predictions across networks and hemispheres, and found that the similarity between each pair of predictions was significantly higher than expected by chance (*q*(FDR) < 0.05, two-sided nonparametric test; Fig. 2d). This demonstrates that each of these cortical networks represents a substantial amount of redundant information (Fig. 2e), and suggests that future brain-computer interfaces could attain good performance even while selectively recording from brain regions that are most accessible or intact.

## Decoder applications and privacy implications

To demonstrate the wide range of potential applications for our decoder, we trained a single semantic language decoder for each subject using brain responses during story perception, and then applied it on brain responses during other tasks.

### Imagined speech decoding

A key task for brain-computer interfaces is decoding covert speech that a subject imagines in the absence of external stimuli^3^. To test whether our language decoder can be used to decode covert speech, subjects imagined telling five one-minute stories while being recorded with fMRI, and separately told the same stories outside of the scanner to provide reference transcripts. Given fMRI recordings for each pair of stories, we correctly identified which recording corresponded to which story by comparing the decoder predictions to the reference transcripts (100% pairwise identification accuracy; Fig. 3a). Across stories, decoder predictions were significantly more similar to their reference transcripts than expected by chance (*p* < 0.05, one-sided nonparametric test). Qualitative analysis shows that the decoder can recover the meaning of imagined stimuli (Fig. 3b; Supplementary Table 2).

**Fig. 3.**
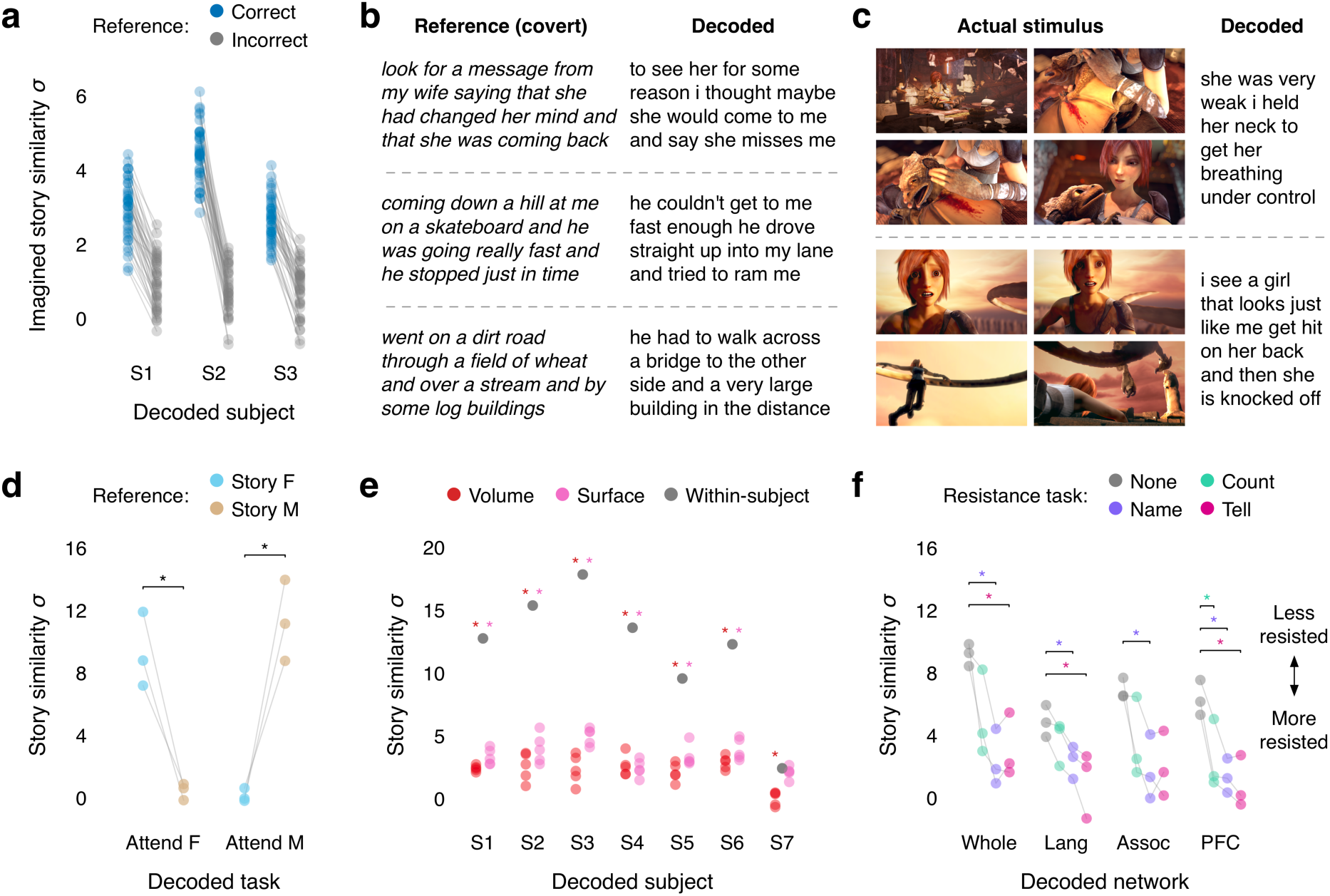
Decoder applications and privacy implications. (**a**) To test whether the language decoder can transfer to imagined speech, subjects imagined telling five 1-minute test stories twice. Single-trial brain responses were decoded and compared to reference transcripts that were separately recorded from the same subjects. For every pair of stories, the correct pairing of predictions and transcripts (prediction 1 with transcript 1, prediction 2 with transcript 2) had higher similarity scores than the incorrect pairing (prediction 1 with transcript 2, prediction 2 with transcript 1). (**b**) Reference transcripts are shown alongside decoder predictions for three imagined stories for one subject. (**c**) To test whether the language decoder can transfer to another modality, subjects watched four silent short films. Single-trial brain responses were decoded using the language decoder. Decoder predictions were significantly related to the films (*q*(FDR) < 0.05, one-sided nonparametric test), and often accurately described film events. Frames from two scenes are shown alongside decoder predictions for one subject. (**d**) To test whether the decoder is modulated by attention, subjects listened to a multi-speaker stimulus that overlays stories told by a female and a male speaker while attending to one or the other. Decoder predictions were significantly more similar to the attended story than to the unattended story (* indicates *q*(FDR) < 0.05 across *n* = 3 subjects, one-sided paired *t*-test; *t*(2) = 6.15 for the female speaker, *t*(2) = 6.45 for the male speaker). Markers indicate individual subjects. (**e**) To test whether decoding can succeed without training data from a particular subject, decoders were trained on brain responses from 5 sets of other subjects (indicated by markers) aligned using volumetric and surface-based methods. Cross-subject decoders performed substantially worse than within-subject decoders (* indicates *q*(FDR) < 0.05, two-sided *t*-test), and barely above chance, suggesting that within-subject training data is critical. (**f**) To test whether decoding can be consciously resisted, subjects performed three resistance strategies: counting, naming animals, and telling a different story. Decoding performance under each condition was compared to a passive listening condition. Naming animals significantly lowered decoding performance in each cortical network (* indicates *q*(FDR) < 0.05 across *n* = 3 subjects, one-sided paired *t*-test; *t*(2) = 6.95 for the whole brain, *t*(2) = 4.93 for the language network, *t*(2) = 3.89 for the association network, *t*(2) = 151.20 for the prefrontal network), demonstrating that decoding can be resisted. Markers indicate individual subjects. For all results, decoding scores depend on the length of the stimuli, so decoding scores are not comparable across different experiments (perceived speech, imagined speech, perceived movie, multi-speaker, decoder resistance).

### Cross-modal decoding

Semantic representations are shared between language and a range of perceptual and conceptual processes^20, 25, 26^, suggesting that, unlike previous language decoders that used mainly motor signals^1, 2^, our semantic language decoder may be able to reconstruct language descriptions from brain responses to non-linguistic tasks. To test this, subjects watched four short films without sound while being recorded with fMRI, and the recorded responses were decoded using the semantic language decoder. We compared the decoded word sequences to audio descriptions of the films for the visually impaired (see Methods), and found that they were significantly more similar than expected by chance (*q*(FDR) < 0.05, one-sided nonparametric test). Qualitatively, the decoded sequences accurately described events from the films (Fig. 3c; Supplementary Table 3). This suggests that a single semantic decoder trained during story perception could be used to decode a range of semantic tasks.

### Attention decoding

Since semantic representations are modulated by attention^27^, our decoder should selectively reconstruct attended stimuli^28, 29^. Beyond improving decoding performance in complex and noisy environments, this could enable our decoder to monitor attention levels and notify subjects when they start to get distracted. To test this, subjects listened to two repeats of a multi-speaker stimulus that was constructed by temporally overlaying a pair of stories told by female and male speakers, while attending to a different speaker for each repeat. Decoder predictions were significantly more similar to the attended story than to the unattended story (*q*(FDR) < 0.05 across subjects, one-sided paired *t*-test), demonstrating that the language decoder selectively reconstructs attended stimuli (Fig. 3d).

### Privacy implications

An important ethical consideration for semantic decoding is its potential to compromise mental privacy^9^. To test if decoders can be trained without a person’s cooperation, we attempted to decode perceived speech from each subject using decoders trained on data from other subjects. For this analysis, we collected data from seven subjects as they listened to five hours of narrative stories. These data were anatomically aligned across subjects using volumetric and surface-based methods (see Methods). Decoders trained on cross-subject data performed barely above chance, and significantly worse than decoders trained on within-subject data (*q*(FDR) < 0.05, two-sided *t*-test). This suggests that subject cooperation remains necessary for decoder training (Fig. 3e; Supplementary Table 4).

To test if a decoder trained with a person’s cooperation can later be consciously resisted, subjects performed three covert cognitive tasks—calculation (“count by sevens”), semantic memory (“name and imagine animals”), and imagined speech (“tell a different story”)—while listening to segments from a narrative story. We found that performing the semantic memory task significantly lowered decoding performance relative to a passive listening baseline for each cortical network (*q*(FDR) < 0.05 across subjects, one-sided paired *t*-test), demonstrating that semantic decoding can be consciously resisted (Fig. 3f).

## Sources of decoding error

To identify potential avenues for improvement, we assessed whether decoding error during story perception reflects random noise in the brain recordings, model misspecification, or both (Fig. 4a).

**Fig. 4.**
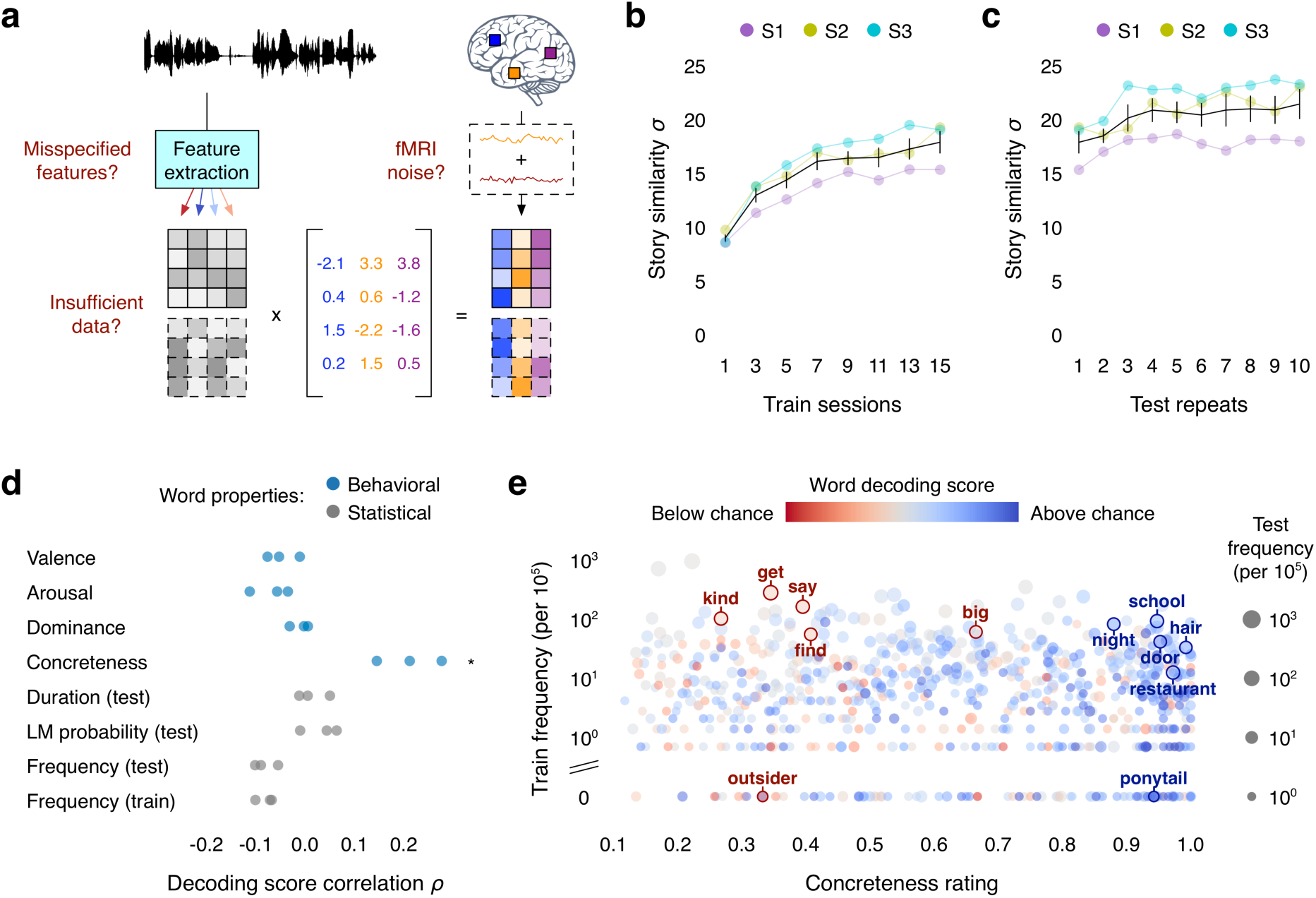
Sources of decoding error. (**a**) Potential factors limiting decoding performance. (**b**) To test if decoding performance is limited by the size of the training dataset, decoders were trained on different amounts of data. Decoding performance increased with the amount of training data collected from each subject but plateaued after 7 scanning sessions (7.5 h) and did not substantially increase up to 15 sessions (16 h). (**c**) To test if decoding performance is limited by noise in the test data, the signal-to-noise ratio of the test responses was artificially raised by averaging across repeats of the test story. Decoding performance slightly increased with the number of averaged responses. (**d**) To test if decoding performance is limited by model misspecification, word-level decoding scores were compared to behavioral ratings and dataset statistics (* indicates *q*(FDR) < 0.05 for all subjects, two-sided permutation test). Markers indicate individual subjects. (**e**) Decoding performance was significantly correlated with word concreteness—suggesting that model misspecification contributes to decoding error—but not word frequency in the training stimuli—suggesting that model misspecification is not caused by noise in the training data. For all results, black lines indicate the mean across subjects and error bars indicate standard error of the mean (*n* = 3).

BOLD fMRI recordings typically have a low signal-to-noise ratio (SNR). During model estimation, the effects of noise in the training data can be reduced by increasing the size of the dataset. To evaluate if decoding performance is limited by the size of our training dataset, we trained decoders using different amounts of data. While decoding performance increased with the amount of training data, most of the improvement occurred by the seventh scanning session—or 7.5 hours—suggesting that simply collecting more data may not substantially improve decoding performance (Fig. 4b).

Low SNR in the test data may also limit the amount of information that can be decoded. To evaluate whether future improvements to single-trial fMRI SNR might improve decoding performance, we artificially increased SNR by averaging brain responses collected during different repeats of the test story. Decoding performance slightly increased with the number of averaged responses (Fig. 4c), suggesting that some component of the decoding error reflects noise in the test data.

Finally, to evaluate if decoding performance is limited by model misspecification—such as using suboptimal features to represent language stimuli—we tested whether the decoding error follows systematic patterns. We scored how well each individual word was decoded across six test stories (see Methods) and compared the scores to behavioral word ratings and dataset statistics. If the decoding error were solely caused by noise in the test data, all words should be equally affected. However, we found that decoding performance was significantly correlated with behavioral ratings of word concreteness (rank correlation *ρ* = 0.15-0.28, *q*(FDR) < 0.05), suggesting that the decoder is worse at recovering words with certain semantic properties (Fig. 4d). Notably, decoding performance was not significantly correlated with word frequency in the training stimuli, suggesting that model misspecification is not primarily caused by noise in the training data (Fig. 4e).

Our results indicate that model misspecification is a major source of decoding error separate from random noise in the training and test data. We expect computational advances—such as the development of better models for extracting semantic features—to substantially reduce model misspecification and improve decoding performance.

## Discussion

This study demonstrates that the meaning of perceived and imagined stimuli can be decoded from BOLD fMRI recordings into continuous language, marking an important step for non-invasive brain-computer interfaces. While existing invasive methods decode from motor representations of articulator and hand motion^1–3^, our non-invasive method decodes directly from semantic representations of language meaning. Our analyses also suggest that semantic decoders trained during story perception can be used to decode other semantic tasks, which may enable augmentative applications such as brain-computer interfaces that transcribe visual experience or translate covert speech into different languages.

While our decoder successfully reconstructs the meaning of language stimuli, it often fails to recover exact words (WER 0.92-0.94). This high WER for novel stimuli is comparable to out-of-set performance for previous invasive decoders^30^, indicating that this loss of specificity is not unique to non-invasive decoding. In our decoder, loss of specificity may occur because different word sequences with similar meanings can evoke similar semantic representations. Future decoders may benefit from modeling language stimuli using a combination of semantic features and lower level features such as phonemes or acoustics.

One other important factor that may improve decoding performance is subject feedback. Previous invasive studies have employed a closed-loop decoding paradigm, where decoder predictions are shown to the subject in real time^2, 3^. This feedback allows the subject to adapt to the decoder, providing them more control over decoder output^31^. While fMRI has lower temporal resolution than invasive methods, closed-loop decoding may still provide many benefits for imagined speech decoding.

Finally, our privacy analysis suggests that subject cooperation is currently required both to train and use the decoder. However, future developments might enable decoders to bypass these requirements. Moreover, even if decoder predictions are inaccurate without subject cooperation, they could be intentionally misinterpreted for malicious purposes. For these and other unforeseen reasons, it is critical to raise awareness of the risks of brain decoding technology and enact policies that protect each person’s mental privacy.

## Methods

### Subjects

Data were collected from three female subjects and four male subjects: S1 (female, age 26 at time of most recent scan), S2 (male, age 36), S3 (male, age 23), S4 (female, age 23), S5 (female, age 23), S6 (male, age 25), and S7 (male, age 24). Data from S1, S2, and S3 were used for the main decoding analyses. Data from all subjects were used to estimate cross-subject decoders for a privacy analysis (Fig. 3e). All subjects were healthy and had normal hearing, and normal or corrected-to-normal vision. The experimental protocol was approved by the Institutional Review Board at the University of Texas at Austin. Written informed consent was obtained from all subjects. To stabilize head motion, subjects wore a personalized head case that precisely fit the shape of each subject’s head.

### MRI data collection

MRI data were collected on a 3T Siemens Skyra scanner at the UT Austin Biomedical Imaging Center using a 64-channel Siemens volume coil. Functional scans were collected using gradient echo EPI with repetition time (TR) = 2.00 s, echo time (TE) = 30.8 ms, flip angle = 71°, multi-band factor (simultaneous multi-slice) = 2, voxel size = 2.6mm x 2.6mm x 2.6mm (slice thickness = 2.6mm), matrix size = (84, 84), and field of view = 220 mm.

Anatomical data for all subjects except S2 were collected using a T1-weighted multi-echo MP-RAGE sequence on the same 3T scanner with voxel size = 1mm x 1mm x 1mm following the Freesurfer morphometry protocol. Anatomical data for subject S2 were collected on a 3T Siemens TIM Trio scanner at the UC Berkeley Brain Imaging Center with a 32-channel Siemens volume coil using the same sequence.

### Cortical networks

Whole brain MRI data were partitioned into 3 cortical networks: the classical language network, the parietal-temporal-occipital association network, and the prefrontal network.

The classical language network was separately localized in each subject using an auditory localizer and a motor localizer. Auditory localizer data were collected in one 10 min scan. The subject listened to 10 repeats of a 1 min auditory stimulus containing 20 s of music (Arcade Fire), speech (Ira Glass, *This American Life*), and natural sound (a babbling brook). To determine whether a voxel was responsive to auditory stimulus, the repeatability of the voxel response across the 10 repeats was calculated using an *F* statistic. This map was used to define the auditory cortex (AC). Motor localizer data were collected in two identical 10 min scans. The subject was cued to perform six different tasks (“hand”, “foot”, “mouth”, “speak”, “saccade”, and “rest”) in a random order in 20 s blocks. For the “speak” cue, subjects were instructed to self-generate a narrative without vocalization. The weight map for the “speak” cue was used to define Broca’s area and the superior ventral premotor (sPMv) speech area.

The parietal-temporal-occipital association network and the prefrontal network were localized in each subject using Freesurfer anatomical ROIs. Voxels identified as part of the classical language network (AC, Broca’s area, and sPMv) were excluded from the parietal-temporal-occipital association and prefrontal networks.

### Experimental tasks

The model training dataset consisted of 82 5-15 min stories taken from *The Moth Radio Hour* and *Modern Love*. In each story, a single speaker tells an autobiographical narrative. Each story was played during a separate fMRI scan with a buffer of 10 s of silence before and after the story. These data were collected during 16 scanning sessions, with the first session consisting of the anatomical scan and localizers, and the 15 subsequent sessions each consisting of 5 or 6 stories. All 15 story sessions were collected for subjects S1, S2, and S3. The first 5 story sessions were collected for the remaining subjects.

Stories were played over Sensimetrics S14 in-ear piezoelectric headphones. The audio for each stimulus was filtered to correct for frequency response and phase errors induced by the headphones using calibration data provided by Sensimetrics and custom Python code (https://github.com/alexhuth/sensimetrics_filter). All stimuli were played at 44.1 kHz using the pygame library in Python.

Each story was manually transcribed by one listener. Certain sounds (for example, laughter and breathing) were also marked to improve the accuracy of the automated alignment. The audio of each story was then downsampled to 11kHz and the Penn Phonetics Lab Forced Aligner (P2FA)^32^ was used to automatically align the audio to the transcript. After automatic alignment was complete, Praat^33^ was used to check and correct each aligned transcript manually.

The model testing dataset consisted of five different fMRI experiments: perceived speech, imagined speech, perceived movie, multi-speaker, and decoder resistance. In the perceived speech experiment, subjects listened to 5-15 min stories from *The Moth Radio Hour*, *Modern Love*, and *The Anthropocene Reviewed*. These test stories were held out from model training. Each story was played during a single fMRI scan with a buffer of 10 s of silence before and after the story.

In the imagined speech experiment, subjects imagined telling 1 min segments from five *Modern Love* stories that were held out from model training. Subjects learned an ID associated with each segment (“alpha”, “bravo”, “charlie”, “delta”, “echo”). Subjects were cued with each ID over headphones and imagined telling the corresponding segment from memory. Each story segment was cued twice in a single 14 min fMRI scan, with 10 s of preparation time after each cue and 10 s of rest time after each segment.

In the perceived movie experiment, subjects viewed four 4-6 min movie clips from animated short films: “La Luna” (Pixar Animation Studios), “Presto” (Pixar Animation Studios), “Partly Cloudy” (Pixar Animation Studios), and “Sintel” (Blender Foundation). The movie clips were self-contained and almost entirely devoid of language. The original high-definition movie clips were cropped and downsampled to 727 x 409 pixels. Subjects were instructed to pay attention to the movie events. Notably, subjects were not instructed to generate an internal narrative. Each movie clip was presented without sound during a single fMRI scan, with a 10 s black screen buffer before and after the movie clip.

In the multi-speaker experiment, subjects listened to two repeats of a 6 min stimulus constructed by temporally overlaying a pair of stories from *The Moth Radio Hour* told by a female and a male speaker. Both stories were held out from model training. Subjects attended to the female speaker for one repeat and the male speaker for the other, with the order counterbalanced across subjects. Each repeat was played during a single fMRI scan with a buffer of 10 s of silence before and after the stimulus.

In each trial of the decoder resistance experiment, subjects were played one of four 80 s segments from a test story over headphones. Before the segment, subjects were cued to perform one of four cognitive tasks (“listen”, “count”, “name”, “tell”). For the “listen” cue, subjects passively listened to the story segment. For the “count” cue, subjects counted by sevens in their heads. For the “name” cue, subjects named and imagined animals in their heads. For the “tell” cue, subjects told different stories in their heads. Trials were balanced such that 1) each task was the first to be cued for some segment and 2) each task was cued exactly once for each segment, resulting in a total of 16 trials. We conducted two 14 min fMRI scans each comprising 8 trials, with 10 s of preparation time after each cue and 10 s of rest time after each trial.

### Language model

Generative Pre-trained Transformer (GPT) is a 12 layer neural network which uses multi-head self-attention to combine representations of each word in a sequence with representations of previous words^18^. GPT was trained on a large corpus of books to predict the probability distribution over the next word 𝑠_𝑛_ in a sequence (𝑠_1_, 𝑠_2_, . . ., 𝑠_𝑛-1_).

We fine-tuned GPT on a corpus comprising Reddit comments (over 200 million total words) and 240 autobiographical stories from *The Moth Radio Hour* and *Modern Love* that were not used for decoder training or testing (over 400,000 total words). The model was trained for 50 epochs with a maximum context length of 100.

GPT estimates a prior probability distribution 𝑃(𝑆) over word sequences. Given a word sequence 𝑆 = (𝑠_1_, 𝑠_2_, . . ., 𝑠_𝑛_), GPT computes the probability of observing 𝑆 in natural language by multiplying the probabilities of each word conditioned on the previous words:  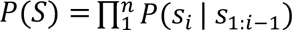 where 𝑠_1:0_ is the empty sequence ∅.

GPT is also used to extract semantic features from language stimuli. In order to successfully perform the next word prediction task, GPT learns to extract quantitative features that capture the meaning of input sequences. Given a word sequence 𝑆 = (𝑠_1_, 𝑠_2_, . . ., 𝑠_𝑛_), the GPT hidden layer activations provide vector embeddings that represent the meaning of the most recent word 𝑠_!_ in context. Previous studies have shown that middle layers of language models extract the best semantic features for predicting brain responses to natural language^13, 14, 16, 17^.

### Encoding model

In voxel-wise modeling, quantitative features are extracted from stimulus words, and regularized linear regression is used to estimate a set of weights that predict how each feature affects the BOLD signal in each voxel.

A stimulus matrix was constructed from the training stories. For each word-time pair (𝑠_𝑖_, 𝑡_𝑖_) in each story, we provided the word sequence (𝑠_𝑖_, 𝑠_𝑖-4_, . . ., 𝑠_𝑖-1_, 𝑠_𝑖_) to the GPT language model and extracted semantic features of 𝑠_𝑖_ from the ninth layer. This yields a new list of vector-time pairs (𝑀_𝑖_, 𝑡_𝑖_) where 𝑀_𝑖_ is a 768-dimensional semantic embedding for 𝑠_𝑖_. These vectors were then resampled at times corresponding to the fMRI acquisitions using a 3-lobe Lanczos filter.

A linearized finite impulse response (FIR) model was fit to every cortical voxel in each subject’s brain. A separate linear temporal filter with four delays (𝑡 − 1, 𝑡 − 2, 𝑡 − 3, and 𝑡 − 4 time-points) was fit for each of the 768 features, yielding a total of 3,072 features. With a TR of 2 s this was accomplished by concatenating the feature vectors from 2, 4, 6, and 8 s earlier to predict responses at time 𝑡. Taking the dot product of this concatenated feature space with a set of linear weights is functionally equivalent to convolving the original stimulus vectors with linear temporal kernels that have non-zero entries for 1-, 2-, 3-, and 4-time-point delays. Before doing regression, we first z-scored each feature channel across the training matrix. This was done to match the features to the fMRI responses, which were z-scored within each scan.

The 3,072 weights for each voxel were estimated using L2-regularized linear regression (also known as ridge regression). The regression procedure has a single free parameter which controls the degree of regularization. This regularization coefficient is found for each voxel in each subject by repeating a regression and cross-validation procedure 50 times. In each iteration, approximately a fifth of the time-points were removed from the model training dataset and reserved for validation. Then the model weights were estimated on the remaining time-points for each of 10 possible regularization coefficients (log spaced between 10 and 1,000). These weights were used to predict responses for the reserved time-points, and then *R*^2^ was computed between actual and predicted responses. For each voxel, the regularization coefficient is chosen as the value that led to the best performance, averaged across bootstraps, on the reserved time-points. The top 10,000 cortical voxels based on cross-validation performance were used for decoding.

The encoding model estimates a function 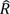 that maps from semantic features 𝑆 to predicted brain responses 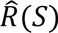. Assuming that BOLD signals are affected by Gaussian additive noise, recorded brain responses 𝑅 and semantic features 𝑆 can be jointly modeled as a multivariate Gaussian distribution (𝑅, 𝑆)^12^. The likelihood of observing responses 𝑅 given semantic features 𝑆 can then be expressed as a multivariate Gaussian distribution 𝑃(𝑅 | 𝑆) with mean 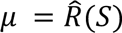 and covariance 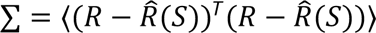. The noise covariance ∑ was estimated using a bootstrap procedure. Each story was held out from the model training dataset, and an encoding model was estimated using the remaining data. A bootstrap noise covariance matrix for the held out story was computed using the residuals between the predicted responses and the true responses. We estimated ∑ by averaging the bootstrap noise covariance matrices across held out stories.

All model fitting and analysis was performed using custom software written in Python, making heavy use of NumPy^34^, SciPy^35^, PyTorch^36^, HuggingFace^37^, and pycortex^38^.

### Word rate model

A word rate model was estimated for each subject to predict when words were perceived or imagined. The word rate at each fMRI acquisition was defined as the number of stimulus words that occurred since the previous acquisition. Regularized linear regression is used to estimate a set of weights that predict the word rate 𝑤 from the brain responses 𝑅. Brain responses were restricted to the classical language network. A separate linear temporal filter with four delays (𝑡 + 1, 𝑡 + 2, 𝑡 + 3, and 𝑡 + 4) was fit for each voxel. With a TR of 2 s this was accomplished by concatenating the responses from 2, 4, 6, and 8 s later to predict the word rate at time *t*. Given novel brain responses, this model predicts the word rate at each acquisition. The time between consecutive acquisitions (2 s) is then evenly divided by the predicted word rates (rounded to the nearest nonnegative integers) to predict word times.

### Beam search decoder

Under Bayes’ theorem, the distribution 𝑃(𝑆 | 𝑅) over word sequences given brain responses can be factorized into a prior distribution 𝑃(𝑆) over word sequences and an encoding distribution 𝑃(𝑅 | 𝑆) over brain responses given word sequences. Given novel brain responses 𝑅*_test_* the most likely word sequence 𝑆*_test_* that the subject is hearing or imagining could theoretically be identified by evaluating 𝑃(𝑆)—with the language model—and 𝑃(𝑅*_test_* | 𝑆)—with the subject’s encoding model—for all possible word sequences 𝑆. However, the combinatorial structure of natural language makes it computationally infeasible to evaluate all possible word sequences. Instead, we approximated the most likely word sequence using a beam search algorithm.

The decoder maintains a beam containing the 𝑘 most likely word sequences. The beam is initialized with an empty word sequence. When new words are detected by the word rate model, the language model generates continuations for each candidate 𝑆 in the beam. The language model is provided with the words (𝑠_𝑛-1_, . . ., 𝑠_𝑛-1"_) that occur in last 8 seconds of the candidate, and predicts the distribution 𝑃(𝑠_𝑛_ | 𝑠_𝑛-𝑖_, . . ., 𝑠_𝑛-1_) over the next word. Nucleus sampling^39^ is used to identify words that belong to the top 𝑝 percent of the probability mass and have a probability within a factor 𝑟 of the most likely word. Nucleus words that occur in the language model input (𝑠_𝑛-𝑖_, . . ., 𝑠_𝑛-1_) are filtered out, as language models have been shown to be biased towards such words. Each word in the remaining nucleus is appended to the candidate to form a continuation 𝐶.

The encoding model scores each continuation by the likelihood 𝑃(𝑅_test_ | 𝐶) of observing the recorded brain responses. The 𝑘 most likely continuations across all candidates are retained in the beam. To increase the diversity of the beam, we accepted a maximum of 5 continuations for each candidate. To increase linguistic coherence, the number of accepted continuations for a candidate was determined by the probability of that candidate under the language model. Candidates in the top quintile under 𝑃(𝑆) were permitted the maximum 5 continuations. Candidates in the next quintile were permitted 4 continuations, and so on, with candidates in the bottom quintile permitted 1 continuation.

### Decoder parameters

The decoder has several parameters that affect model performance. The encoding model is parameterized by the number of context words provided when extracting GPT embeddings. The noise model is parameterized by a shrinkage factor ɑ that regularizes the covariance ∑. Language model parameters include the length of the input context, the nucleus mass 𝑝 and ratio 𝑟, and the set of possible output words.

All parameters were tuned by grid search and by hand on data collected as subject S3 listened to a calibration story from *The Moth Radio Hour* separate from the training and test stories. We decoded the calibration story using each configuration of parameters. The best performing parameter values were validated and adjusted through qualitative analysis of decoder predictions. The parameters values used in this study provide a default decoder configuration, but in practice can be tuned separately and continually for each subject to improve performance.

To ensure that our results generalize to new subjects and stimuli, we restricted all pilot analyses to data collected as subject S3 listened to the test story “Where There’s Smoke” by Jenifer Hixson from *The Moth Radio Hour*. All pilot analyses on the test story were qualitative. We froze the analysis pipeline before we viewed any results for the remaining subjects, stimuli, and experiments.

### Language similarity metrics

Decoded word sequences were compared to reference word sequences using a range of automated metrics for evaluating language similarity. Word error rate (WER) computes the number of edits (word insertions, deletions, or substitutions) required to change the predicted sequence into the reference sequence. BLEU^40^ computes the number of predicted *n*-grams that occur in the reference sequence (precision). We used the unigram variant BLEU-1. METEOR^41^ combines the number of predicted unigrams that occur in the reference sequence (precision) with the number of reference unigrams that occur in the predicted sequence (recall), and accounts for synonymy and stemming using external databases. BERTScore^42^ uses a bidirectional transformer language model to represent each word in the predicted and reference sequences as a contextualized embedding, and then computes a matching score over the predicted and reference embeddings. We used the recall variant of BERTScore with IDF importance weighting computed across stories in the training dataset. BERTScore was used for all analyses where the language similarity metric is not specified.

For the perceived speech, multi-speaker, and decoder resistance experiments, stimulus transcripts were used as reference sequences. For the imagined speech experiment, subjects told each story segment out loud outside of the scanner, and the audio was recorded and manually transcribed to provide reference sequences. For the perceived movie experiment, official audio descriptions from Pixar Animation Studios were manually transcribed to provide reference sequences for three movies. To compare word sequences decoded from different brain regions (Fig. 2d), each sequence was scored using the other as reference and the scores were averaged (prediction similarity).

We scored the predicted and reference words within a 20 s window around every second of the stimulus (window similarity). Scores were averaged across windows to quantify how well the decoder predicted the full stimulus (story similarity).

To test whether stimulus time-points can be identified using decoder predictions, we performed a post hoc identification analysis using similarity scores between predicted and reference sequences. We constructed a matrix 𝑀 where 𝑀_%/_ reflects the similarity between the 𝑖th predicted window and the 𝑗th reference window. For each time-point 𝑖, we sorted all reference windows by their similarity to 𝑖th predicted window, and scored the time-point by the percentile rank of the 𝑖th reference window. The mean percentile rank for the full stimulus was obtained by averaging percentile ranks across time-points.

### Statistical testing

To test statistical significance of decoding scores, we generated 200 null sequences by sampling from the language model without using any brain data. These sequences reflect the null hypothesis that the decoder does not reconstruct meaningful information about the stimulus from the brain data. We scored the null sequences against the reference sequence to produce a null distribution of decoding scores. We compared the observed decoding scores to this null distribution using a one-sided nonparametric test; *p*-values were computed as the fraction of null sequences with a decoding score greater than or equal to the observed decoding score.

To test statistical significance of the post hoc identification analysis, we randomly shuffled 10-row blocks of the similarity matrix 𝑀 before computing percentile ranks. We evaluated 2,000 shuffles to obtain a null distribution of mean percentile ranks. We compared observed mean percentile rank to this null distribution using a one-sided permutation test; *p*-values were computed as the fraction of shuffles with a mean percentile rank greater than or equal to than the observed mean percentile rank.

Unless otherwise stated, all tests were performed within each subject and then replicated across all subjects (*n* = 7 for the cross-subject decoding analysis shown in Fig. 3e, *n* = 3 for all other analyses). All tests were corrected for multiple comparisons when necessary using the false discovery rate (FDR)^43^.

### Word decoding performance

To quantify word-level decoding performance, we represented words using 300-dimensional GloVe embeddings^44^. We considered a 10 s window centered around each stimulus word. We computed the maximum linear correlation between the stimulus word and the predicted words in the window. Then, for each of the 200 null sequences, we computed the maximum linear correlation between the stimulus word and the null words in the window. The match score for the stimulus word was defined as the number of null sequences with a maximum correlation less than the maximum correlation of the predicted sequence. Match scores above 100 indicate higher decoding performance than expected by chance, while match scores below 100 indicate lower decoding performance than expected by chance.

Match scores were averaged across all occurrences of a word across six test stories. The word-level match scores were compared to behavioral ratings of valence (pleasantness), arousal (intensity of emotion), dominance (degree of exerted control), and concreteness (degree of sensory or motor experience)^45, 46^. Each set of behavioral ratings was linearly rescaled to be between 0 and 1. The word-level match scores were also compared to word duration in the test dataset, language model probability in the test dataset, word frequency in the test dataset, and word frequency in the training dataset.

### Anatomical alignment

To test if decoders could be estimated without any training data from a target subject, volumetric^47^ and surface-based^48^ methods were used to anatomically align training data from separate source subjects into the volumetric space of the target subject.

For volumetric alignment, we used the *get_mnixfm* function in pycortex^38^ to compute a linear map from the volumetric space of each source subject to the MNI template space. This map was applied to recorded brain responses for each training story using the *transform_to_mni* function in pycortex. We then used the *transform_mni_to_subject* function in pycortex to map the responses in MNI152 space to the volumetric space of the target subject. We z-scored the response time-course for each voxel in the volumetric space of the target subject.

For surface-based alignment, we used the *get_mri_surf2surf_matrix* function in pycortex to compute a map from the surface vertices of each source subject to the surface vertices of the target subject. This map was applied to the recorded brain responses for each training story. We then mapped the surface vertices of the target subject into the volumetric space of the target subject using the *line-nearest* scheme in pycortex. We z-scored the response time-course for each voxel in the volumetric space of the target subject.

Each source subject independently produces a set of aligned responses for the target subject. To aggregate across source subjects, we used a bootstrap procedure to sample five sets of source subjects. For each bootstrap, the aligned responses were averaged across the sampled source subjects. To estimate the noise model ∑, aligned responses from a single, randomly sampled source subject were used to compute the bootstrap noise covariance matrix for each training story. Separate encoding models were estimated using volumetric and surface-based alignment. Cross-subject decoders were evaluated by decoding actual responses recorded from the target subject.

## Data availability

All data used in the analysis will be made publicly available at OpenNeuro before publication.

## Code availability

All code used in the analysis will be made publicly available at GitHub before publication.

## Acknowledgements

We thank J. Wang, XX. Wei, and L. Hamilton for comments on the manuscript. This work was supported by the Whitehall Foundation, Alfred P. Sloan Foundation, and the Burroughs Wellcome Fund.

## Author Contributions

Conceptualization J.T. and A.G.H.; Methodology J.T.; Software and resources J.T. and S.J.; Investigation and data curation J.T. and A.L.; Formal analysis and visualization J.T.; Writing (original draft) J.T.; Writing (review and editing) J.T., A.L., S.J., and A.G.H.; Supervision A.G.H.

## Additional Information

Supplementary Information is available for this paper.

Correspondence and requests for materials should be addressed to.

---

**Extended Data Fig. 1.**
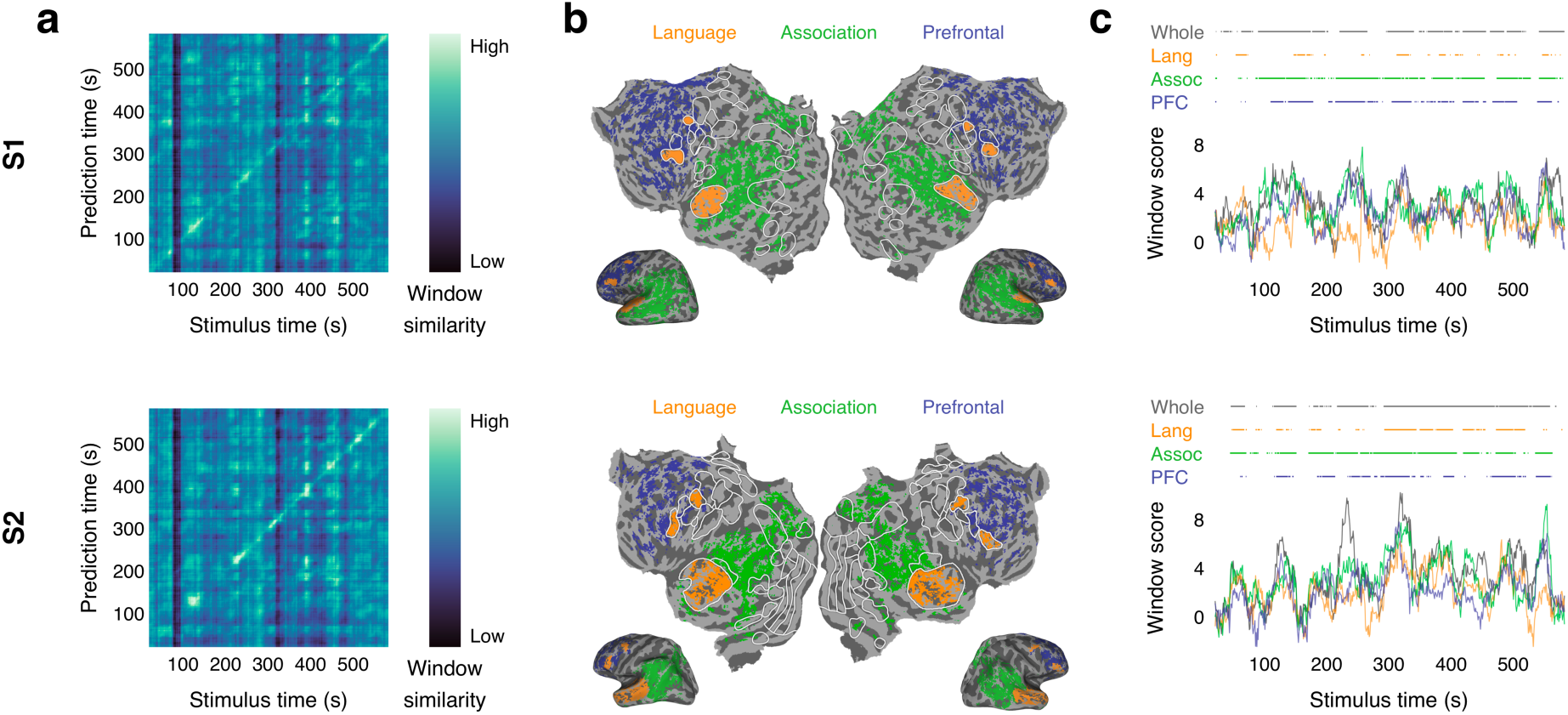
Perceived speech decoding performance. Language decoders were trained for subjects S1 and S2 and then evaluated on single-trial BOLD fMRI responses recorded while the subjects listened to the test story “Where There’s Smoke” by Jenifer Hixson from *The Moth Radio Hour*. (**a**) Identification accuracy for whole brain decoder predictions. The brightness at (𝑖, 𝑗) reflects the BERTScore similarity between the 𝑖th second of the decoder prediction and the 𝑗th second of the actual stimulus. Identification accuracy was significantly higher than expected by chance (*p* < 0.05, one-sided permutation test). (**b**) Cortical networks for subjects S1 and S2. Brain data used for decoding (colored voxels) were partitioned into the classical language network, the parietal-temporal-occipital association network, and the prefrontal network (PFC). (**c**) Decoding performance time-course from each network. Horizontal lines indicate when decoder predictions were significantly more similar to the actual stimulus than expected by chance under the BERTScore metric (*q*(FDR) < 0.05, one-sided nonparametric test). Corresponding results for subject S3 are shown in Figures 1f, 2a, and 2c in the main text.

## Supplementary Information

**Supplementary Table 1.**
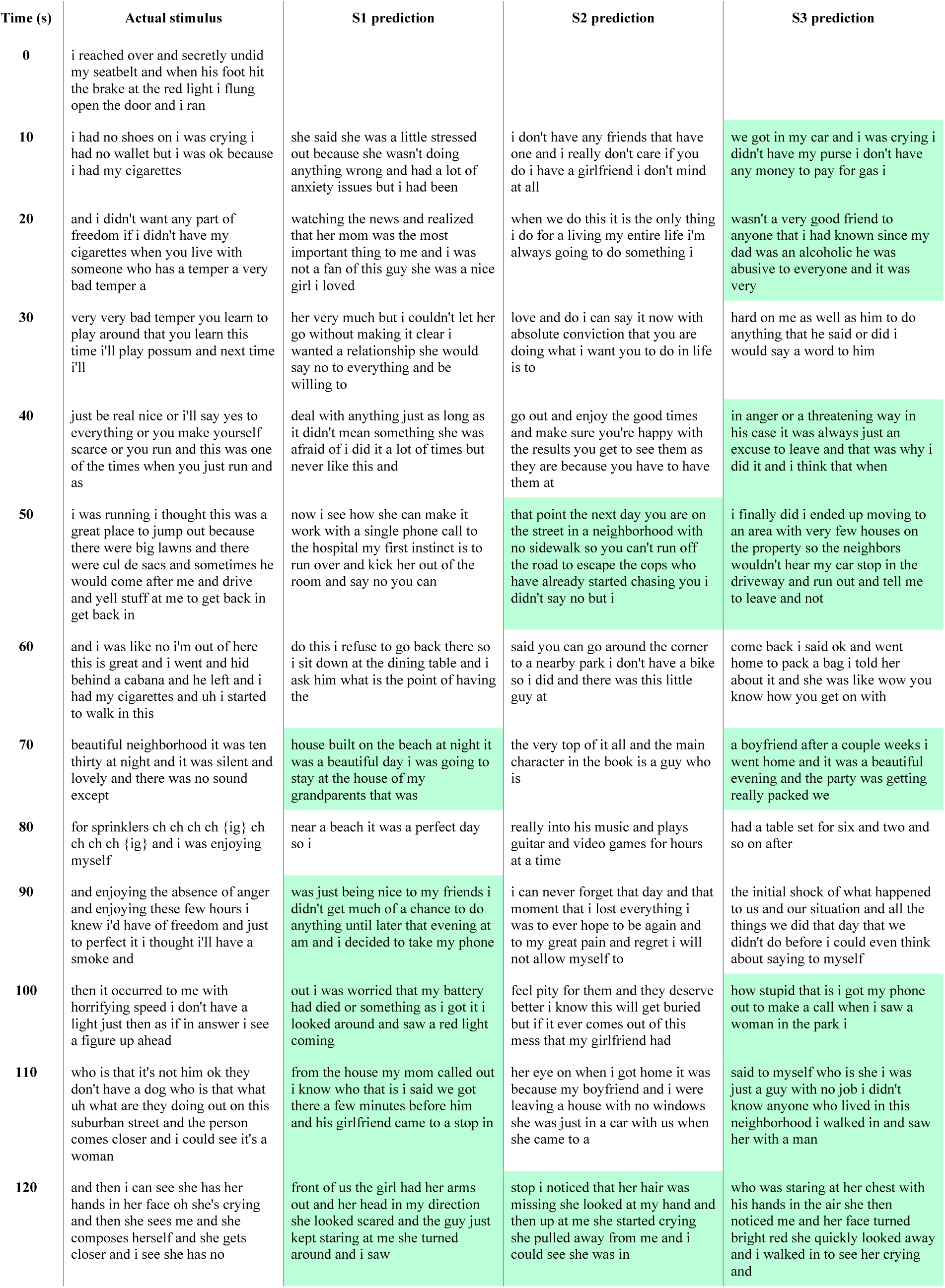
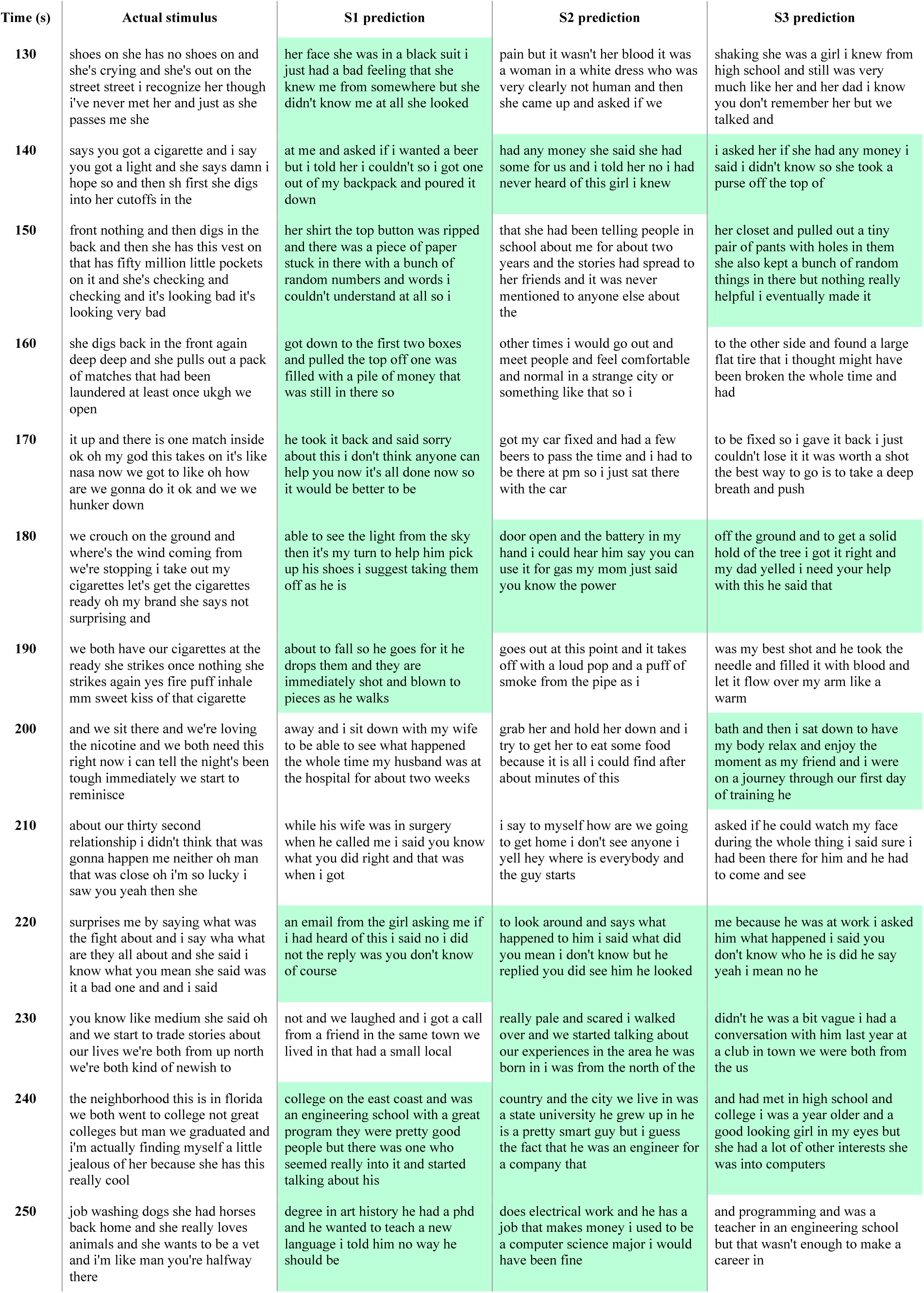
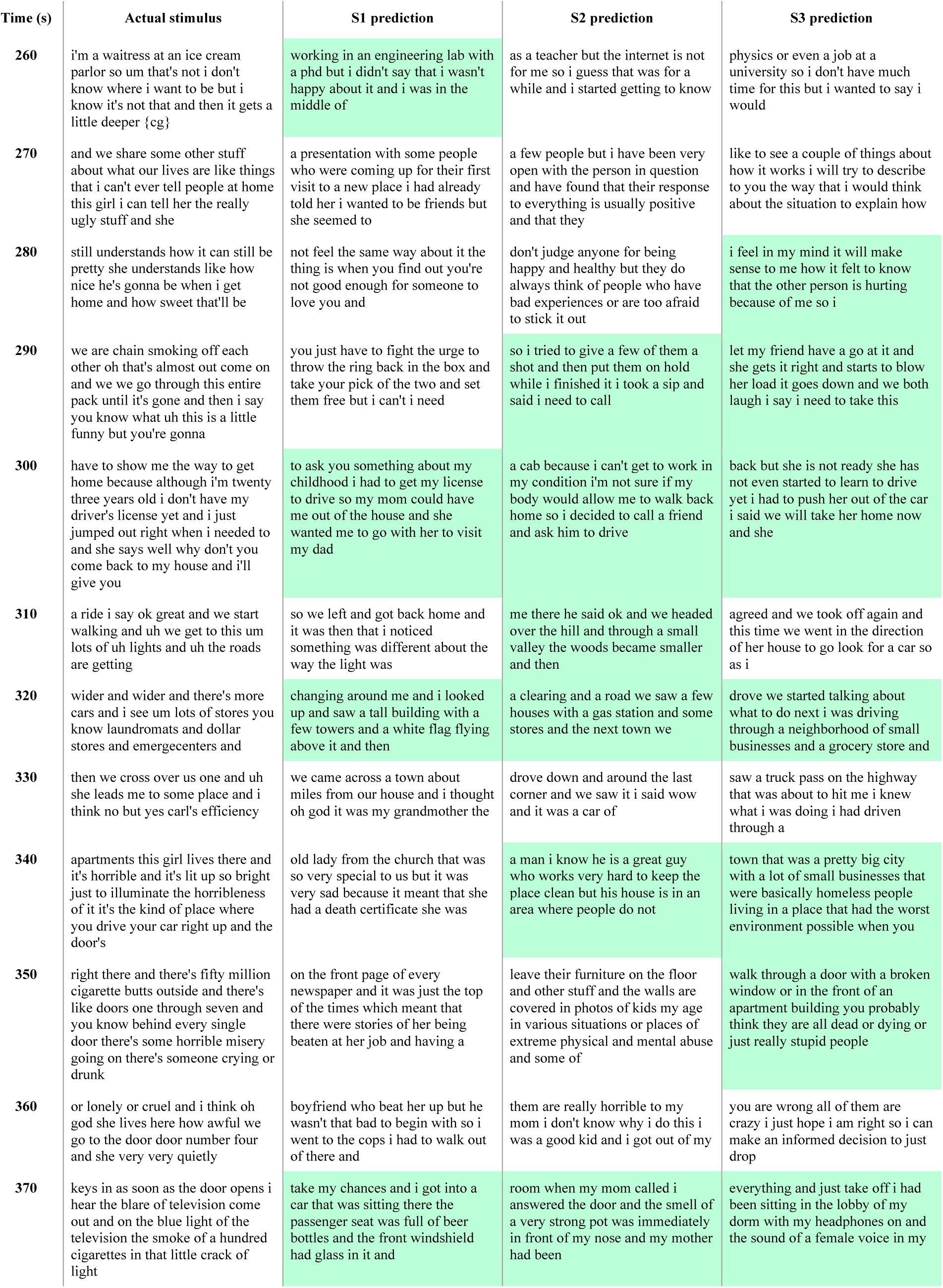
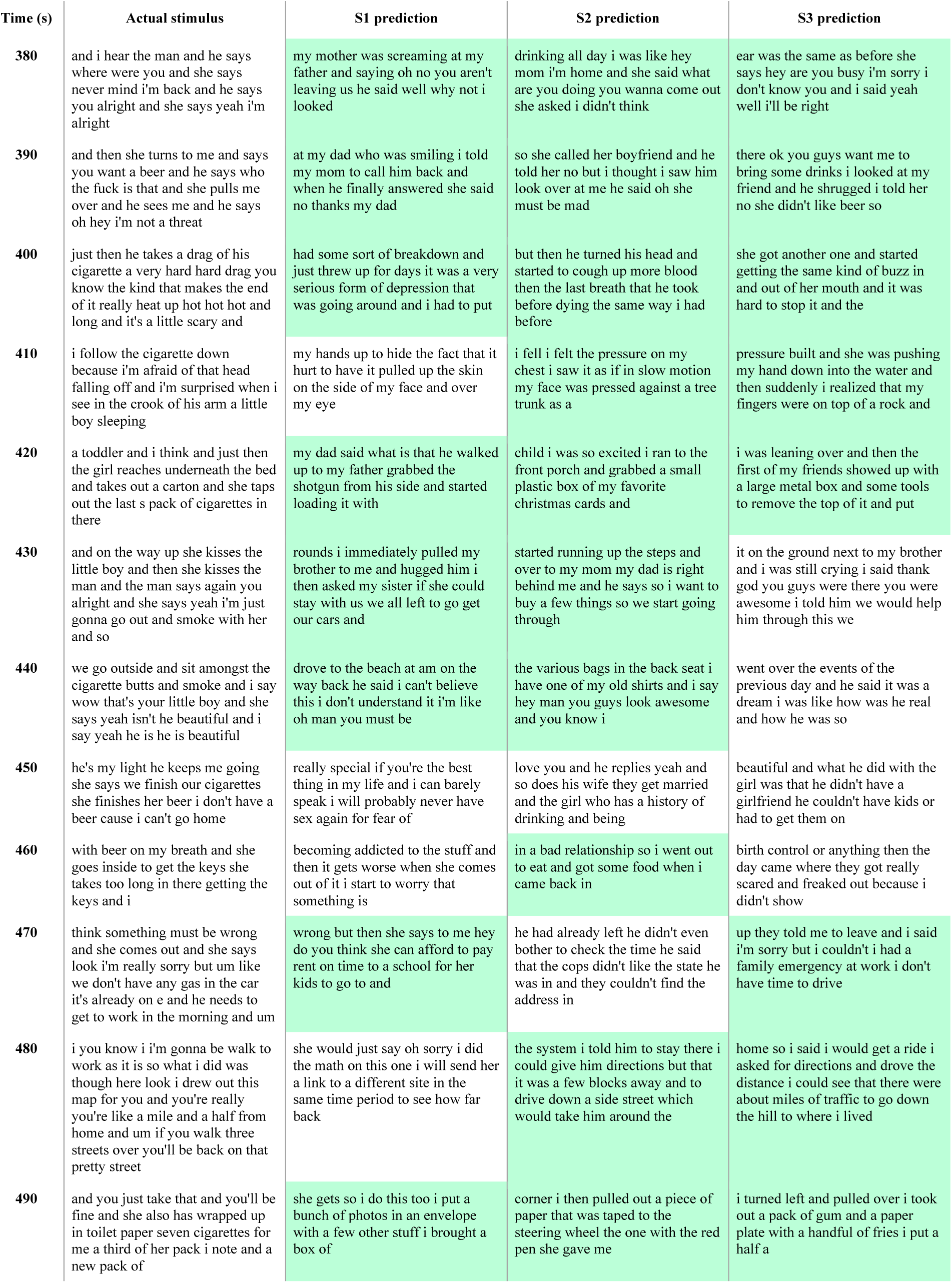
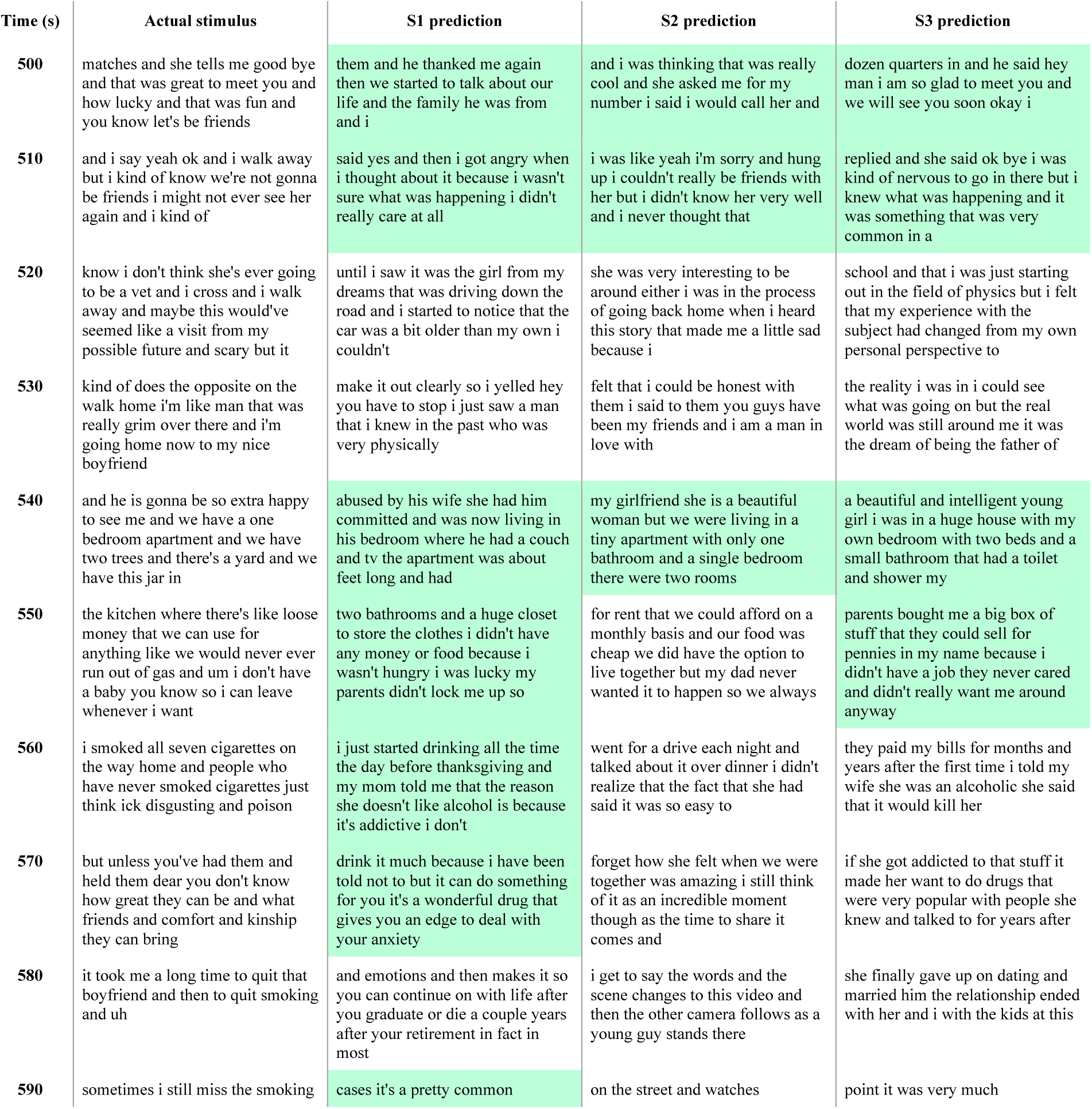
Decoder predictions for a perceived story. Language decoders were trained for each subject and then evaluated on single-trial BOLD fMRI responses recorded while that subject listened to the test story “Where There’s Smoke” by Jenifer Hixson from *The Moth Radio Hour*. The actual stimulus words are shown alongside the decoder predictions for each subject. Highlighted segments were significantly more similar to the actual stimulus than expected by chance under the BERTScore metric (q(FDR) < 0.05, one-sided nonparametric test). The decoders recovered the meaning of the perceived story.

**Supplementary Table 2.** Decoder predictions for imagined stories. Language decoders were trained for each subject (using data collected during story perception) and then evaluated on single-trial BOLD fMRI responses recorded while that subject imagined telling 1-minute segments from five different test stories from *Modern Love*. The story segments that the subjects were instructed to imagine are shown alongside the decoder predictions for each subject. Highlighted segments were significantly more similar to the reference stimulus than expected by chance under the BERTScore metric (q(FDR) < 0.05, one-sided nonparametric test). Decoder predictions were also significantly similar to transcripts recorded from each subject (q(FDR) < 0.05; not shown). Results are shown for the second repeats of each segment; results for the first repeats were similar. The decoders recovered the meaning of the imagined stories.

| Time (s) | Imagined stimulus (Alpha) | S1 prediction | S2 prediction | S3 prediction |
| --- | --- | --- | --- | --- |
| 0 | every morning i woke up with two giant animals nuzzling me panting in my face demanding to be walked i would walk them | he was not happy with the idea of me being on his couch and sleeping with him and then when he had the chance to leave me in | it would be easier to just throw her over my shoulder and leave as she is already dressed and ready for bed when she calls my phone at | we need to know who it is and if they are coming to pick up my girl i will go and call her right away because i want to see her for |
| 10 | and upon my return i'd check the phone hoping i had somehow missed my wife's call saying that she had changed her mind and was moving back she never did | the hall and i saw his face when i said something to him that he couldn't remember and he asked me to come in he didn't really know | am to say that the police are there to talk to me about what happened she was right but not the end result i just had to move on | some reason i thought maybe she would come to me and say she misses me but no she was dead when i found her this is the truth i was in |
| 20 | i tried to carry on i bought groceries said hello to the neighbors i can see now that i was in | me i had a bad experience with drunk drivers at my college and that was all it was the worst part about it i don't | i started going to school and the people around me would say i was depressed and not happy it was like the world was ending i had | the hospital and my parents were at my house we had a party with my family and the doctors and nurses and everything but |
| 30 | shock pretending everything would be ok the house deteriorated the laundry hamper overflowed | remember anything about the accident the car was spinning in the air and the windshield was cracked and i was hanging out of it the front | been living on a diet and on the side of a toilet and a bottle of water and the bottom of my | we got through it just fine because of that we don't have insurance i had an accident on my bike with a tire iron and a |
| 40 | bags of trash clustered in the kitchen my bed was a sleeping bag dog hair accumulated in giant clumps in the corner of rooms and in | passenger side was just a piece of broken metal and a few others were scattered everywhere and the back door was open i knew the next | shoe on the ground my socks are covered in the sand and mud of the driveway the only time i have to | broken back in the process my father died and i had to leave to help him his funeral was a lot like what you hear on television people |
| 50 | filthy patches on the rug i was commuting an hour each way for work and unmoored from my previous life couldn't manage my responsibilities | step was to find a doctor and get help i started to cry and run i ran home i stayed up late until my alarm went off and | drive from one point to another is to the last stop i make in town and to my surprise i don't have a ticket for that one | are all over it all of them but not everyone is interested in you it gets a little confusing to be a single person |

| Time (s) | Imagined stimulus (Bravo) | S1 prediction | S2 prediction | S3 prediction |
| --- | --- | --- | --- | --- |
| 0 | when my son jeff was little he was a pain in the neck about eating on one drive to huntsville alabama he | they got to the point where they were not in any condition to drive and were having a panic attack when i asked her why they | it is my favorite place to spend an evening because of the music it was playing during the concert that made me want to scream in the night at how stupid that | it is a very serious matter i am very concerned with how much time you will spend in jail and i do mean a month and a half ago i would say that |
| 10 | sobbed for seventy minutes i know because i timed it about how we were starving him to death we stopped at a diner and ordered him a meal and he | had to call her in to help out she just looked at me with those eyes and said please do not leave my husband is very upset | song was i thought about telling her but i decided to wait i went home and got some more weed i didn't think she was going to drink i just | if you get into the car and drive to a restaurant in the middle of nowhere and order a beer that is where the food is we decide to take turns with |
| 20 | proceeded to eat about four bites before claiming he was full you might think i would lose my temper but this had happened before so i was prepared | i think he wants to tell me something but he isn't sure i will agree or even respond in kind so he starts with me having sex with him but | felt bad and hoped she wouldn't take it i told her it was okay she took the bottle from me and started drinking it as i held it up | the bill and it comes off as a dollar i then give it to him as i don't want it anymore but it seems like the customer should |
| 30 | with a well planned response i reached over and started eating his food bite by bite i finished everything on his plate figuring | i don't feel anything i say that was wrong and i really wanted it to be true but he wasn't telling the truth the guy who did this was | i had to give her some more because she had already had it but i knew this was going to be hard for her to handle i did this | get a taste of this kind of treatment so i could be on my way i didn't do it all at once or even by |
| 40 | that would teach him to mind his dinner unfortunately the plan had a different effect everywhere we went after that jeff expected me | in prison and we just assumed he had the ability to do that because of how long he lived i got out of there and started moving | to her for over years i know that if she ever had a boyfriend or girlfriend it would be different because i never thought it was possible she had to | accident but the timing was right and it was my choice not to have my father in jail for not getting a job that i could have |
| 50 | to finish his meals it got so i would only order him meals i liked knowing how it would go | around in a new city and then bam they got the phone number and called me at work they had | be very close to a male of my own race so i wasn't as much of a threat to their | used my entire adult life as i had it before my parents had a panic attack when they had to get to work but i still did |

| Time (s) | Imagined stimulus (Charlie) | S1 prediction | S2 prediction | S3 prediction |
| --- | --- | --- | --- | --- |
| 0 | for my third date with tracy i'm taking her to the sunnyside rice shoppe in manhattan you're going to love this place i tell her when i was a kid i used to order pork chow | it was the night before the funeral that i decided to tell my dad he would understand but i was young and in love with the girl at the time we had | i was at a bar and this woman i know who is my friend and who i like a lot tells me to drink this and she | i had a dream about my best friend in college and he told me about a story he had read recently and i was there for him as he was |
| 10 | mai fun and smear it on the plate with ketchup tracy beams at me from the passenger seat she clearly likes that i'm already sharing family stuff | a small plot of land and was building it up in our backyard with some friends she was an incredible cook and always wanted to help us out | puts some in my cup and i feel my face turn red she was laughing and it was a great time so i say excuse me and | sick so i offered him a cup of tea i could smell the sugar and it made me smile because i was already getting better at |
| 20 | i wait for the pedestrians to clear before turning off canal onto mott at bayard i take a left there's the restaurant i say you want to get out and i'll go | with something as a teenager my dad got her into the kitchen and then to the other room so that we could get to work he said we | leave her i get into the car and drive off the main road into a town on the way home and pull in my dad says ok go ahead and | this i didn't think it was fair to take my job too seriously but the boss said i could |
| 30 | park nah says tracy i'll help you find a spot ok | should do this and i told him no i can't and that he should leave we ended up making the deal i don't think we went into | stop my mom asks me to just take a shower i reply oh no that's fine she said i should get some rest i say sure and i head down | go and he was right so i went i spent about minutes looking for the place and came up empty and then i realized that the door was not locked |
| 40 | i drive to the end of bayard but there are no spots at bowery i swing a right then at pell another right | it directly but the first week we did go to a movie a few of us had been to it before but i remember it | stairs and down the hall and turn left and right and then down a couple of floors and finally around another corner and there it | i started to walk away and the guy said hey you want some of my beer i don't need it he says it again i can't |
| 50 | there's what looks like a spot but when we get closer i notice a two pronged fire hydrant protruding from a brick wall nothing on pell i | was really creepy as hell with a giant snake on a string and all it had to do was come and eat me i don't think | is again i was almost sure it was the same one it had a very similar looking body type but i could have been wrong because | really hear you i'm not drunk i get it this time i actually do see it all in slow motion and i can feel the |

| Time (s) | Imagined stimulus (Delta) | S1 prediction | S2 prediction | S3 prediction |
| --- | --- | --- | --- | --- |
| 0 | that spring was promising to be the greatest time of my life i was happily in love for the first time i | they are the same in the real world i was going to tell her about this when i saw the pictures of my wife she looked absolutely beautiful and | they will never be the same after i lost my dad he was my first friend and i knew nothing about him except for how | we were on vacation for a month and were very much in love with the idea of a little girl and a baby she was absolutely beautiful and |
| 10 | saw marko at school and was attracted to his eyes and playful smile one afternoon while walking home from a piano lesson | was sitting with her boyfriend in a movie theater with him we all were on a couch and he was holding her hand while he looked at her pictures i knew | to say it he had this huge tattoo on his arm and a really cute girl who was with him when we walked in and saw his | very independent and when they were gone she would sit up and look around the house and just get upset when i left to go |
| 20 | i saw him coming down the hill on his skateboard he stopped just short of running into me i don't remember us saying | my parents were worried so i asked him how he could tell his story he was very emotional and had to hold himself back i said i have | friend on the ground with a knife he immediately jumped in the car and pulled him away we all sat in silence and i said something to him i think i'm just | to school i ran into a man on his way to work that he was heading towards his car in the middle of the morning and told him what happened and how i had |
| 30 | much we just stood there and smiled but that's all it took to seal the deal and we became inseparable | never been happier and we went to the park and the dog ran out in front of a car and started biting and tearing at it i saw | stressed out from this day forward and that we could discuss it at my parents place or somewhere else for example we had a group of friends who would take | been the victim and what i did not know was that he had taken me to a friend of mine and i was brought to him and the other guys by |
| 40 | one day a group of us decided to climb an abandoned building and jump off its flat roof i was standing on the edge | a man go flying through the air and i thought to myself you want to live that fast but what happens when he lands you only have seconds before the wind rips | a ride down to the river and then drive back up the hill and sit on the rock facing the water while it comes up my friend starts crying he | my boyfriend and his girlfriend he was there with me for most of it i had to wear a shirt in a lot of ways that he hated to see |
| 50 | twelve feet above the dirt my heart racing when marko scooted next to me you are the bravest girl i know he whispered with those words my fear | you apart and then you're gone forever but even then it doesn't last i just went from the best friend who would | runs up to me and hugs me tight and whispers you saved me i just had a thought that was like the one where the person on the other side | so he could look at me like i was ugly he even offered me his hand which i took i felt so small in that moment i just didn't |

**Supplementary Table 2. Decoder predictions for imagined stories.**
| Time (s) | Imagined stimulus (Echo) | S1 prediction | S2 prediction | S3 prediction |
| --- | --- | --- | --- | --- |
| 0 | to reach chimi you take a dirt road then walk through wheat fields across a stream and past timber houses until you climb a small rise and | she was still on the road but in a better part of town we passed by a sign for an old school that had a huge tree | it to be able to walk back home in a straight line or in the direction of a house on a hill without breaking into a run the | he was on the road for about hours and his phone battery was dead we decided to go to my parents house and he went to mine since the |
| 10 | find an unprepossessing building surrounded by the usual troika of himalayan temple attendants dogs scratching at fleas chickens scratching at dirt and monks | house and a little girl riding her bike i said sorry and turned away it was just a game i thought i could get used to i | police report shows a large red and yellow crime scene tape with the name of a local law firm of cops with a black and white id on | one i had at home was just for the kids he was so nice and so gentle but when he left his house that was all there was i could |
| 20 | scratching at well it's not clear their robes cover quite a lot it was a pleasant place but nothing out of the ordinary i gave a prayer | got the car back and i was able to drive to work with my head down i drove by a police station with the license plate of the cop on | their shirt i didn't like it when they were there either so i said well it's too bad because i'm just going to throw a fit i just | do my part he had a small car that i knew he liked so we drove to the restaurant and he opened up the door and took a |
| 30 | wheel a lazy spin and even yawned as i walked into the central chamber in front of us loomed a smoky altar of goddesses at their feet lay offerings of | the side i turned the corner and saw him and the other two cars and one other guy with a baseball bat i was floored the | threw my hands up and walked out i was only at that point and my mom was coming out the door of the room in the corner i just knew it was going to be | deep breath i sat there awkwardly in the passenger seat for the next few minutes with the engine running but after a couple hours i had to |
| 40 | currency garlands of flowers candles fruit the room hummed with flies and noise and tourists but it was also full of an | cops were calling it a hit and run and my brother was still alive they were very respectful and very calm in my family they all thought it | bad and i had to do it so i slowly lifted the cover and saw my favorite piece of cake and chocolate chip ice cream | stop because of the smoke i started coughing and the smell of my own body filled the house it was like an animal attack i was |
| 50 | incredibly concentrated hopefulness something so spirited and real to me that i was brought uncharacteristically to my knees | was ok to talk about it and if you didn't feel comfortable enough you just asked and you know what i felt like i did a | i had just eaten i could see a giant cloud of smoke and heard people shouting and some kind of noise but i just kept walking towards | completely overcome and it wasn't a very healthy and natural thing i couldn't get over the experience of seeing the girl in a hospital bed for |

**Supplementary Table 3.** Decoder predictions for perceived movies. Language decoders were trained for each subject (using data collected during story perception) and then evaluated on single-trial BOLD fMRI responses recorded while that subject watched three animated short films from Pixar Animation Studios (“La Luna”, “Presto”, and “Partly Cloudy”) without sound. Reference transcripts—obtained from official audio descriptions for the visually impaired—are shown alongside the decoder predictions for each subject. Highlighted segments were significantly more similar to the reference words than expected by chance under the BERTScore metric (q(FDR) < 0.05, one-sided nonparametric test). The decoders generated language descriptions that capture the meaning of the perceived movies.

| Time (s) | Description ("La Luna") | S1 prediction | S2 prediction | S3 prediction |
| --- | --- | --- | --- | --- |
| 0 | now over calm waters stars twinkle in the night sky a rowboat glides into view bearing the name la luna as a burly man with |  |  |  |
| 10 | a thick brown mustache works the oars a little boy peeks over the side of the small wooden craft an old man with a long white beard sits near the stern a hanging | it was a beautiful summer day and a pretty girl was standing there and we had to wait at the bottom of a hill for | he is the man of the house and a gentleman in the family the same as your grandmother if i were to meet you on the street and | it was a small thing but i did feel for the man the story of how he died and how the doctors found the |
| 20 | lantern behind him a bird's eye view shows the boat moving steadily across the ocean's dark surface with no land in sight the burly man sets his oars inside | a taxi which wasn't easy considering the nature of the job so i have no idea what they do with the money and it is difficult to imagine the | see you walk away in my mind it would be the exact opposite i was at a bar with two guys and one girl when the first | cause of death the same way as if you had actually committed a crime with that amount of blood i went into a room and put a |
| 30 | the boat and the old man drops an anchor the old man hands the boy a gift he unwraps it to find a brown newsboy hat | company that works with it so many people just walking around the store doing nothing the owner just gives me a smile and says come in and | of them grabbed a glass and filled it with the cheapest beer and asked me to step in he said to me and his two sons if you | blanket over it and left my buddy in there with her and my friend she started crying and he took her in he told me about his little sister who |
| 40 | just like his dad and grandpa's he tries it on the old man turns the boy his way and lifts the hat's brim his father lowers the brim to be like his the | i just like to hang out with these two i was one of them for about a month we both had a history together we would have sex but | can put on a pair of socks i did they were a size too small he gave me a hug and then turned to his younger brother he | was now his wife he had a young daughter and she was very pretty but he kept it down and not made fun of me i still |
| 50 | grandpa tips it up again but the father pushes it back down and sets the boy on the seat next to his grandpa the grandpa turns off the lantern and folds his arms as the old | she never let me go and i just kept saying no she said no i did this i mean it was like the worst thing to have | told them to leave but the youngest one just kept saying no they had no money he was on a bus to get to the next | had the best friends at school but my parents didn't like it anymore so we just went on with our lives my mom and dad |
| 60 | man and his burly son both sit with their arms crossed the boy takes notice and folds his arms as well the burly man wipes his nose | happen to me my brother got drunk and killed himself my parents were in tears my sister and i cried and my dad stayed quiet as | stop and that he had been caught doing something i had done in my life i just smiled and said hi to him as i passed and | didn't really agree with each other and it wasn't a happy situation at all my brother was very angry he started screaming at me |
| 70 | the boy does the same he watches his grandpa twist his finger in his ear he mimics him | if to comfort me he tried to hold back but my brother had his hand on my shoulder he squeezed hard i cried harder then i just went | i got a hug in return i remember my mother crying a little and her hand rubbing my back and the next moment she was | i yelled at him and i hit him with my hand the other one went through my face i heard a crack and my vision started to |
| 80 | as a huge bright moon rises over the watery horizon the wide eyed boy's jaw drops the rippling waters around the rowboat | to sleep i was in a dark room in an abandoned factory where i knew nothing but the shadows it was a few hours later and | in a coma the hospital was a few miles from where i lived in the state that was so horrible it was almost | fade as the first light of dawn showed me a blurry view of a city that was far too bright for the sun i realized that i had |
| 90 | reflect the massive orb's glow the father hands the boy an anchor then opens a compartment the boy goes to drop the anchor his grandpa gestures for him to wait the | the building was dark and cold i got my laptop out and set it up to show her that i was typing it and when she asked how | a ghost town after the fire we were out of gas and i got my friend to help me load my truck he said to do it right away so he | traveled for hours to the airport and i was very hungry i went inside to grab something and the manager said i need to take the phone back to my seat he explained that we had |
| 100 | burly man lifts a long ladder from the compartment the boy watches the seemingly endless rungs pass by his father turns to him he gestures to head up | it was i explained how she could see the page from a computer that was not there before i left her office and said good luck to | took off the second time i had to lift it i got it about feet up and he grabbed it and pulled me up after him so | to stop in front of the theater so the car would park right next to it we then sat down and talked about this girl we |
| 110 | his grandpa also gestures for the boy to go on the boy slings the anchor over his shoulder ties its rope around his chest then heads up the ladder about | her she asked me to wait outside i took my hat off and held it under my arm so as not to fall and fell i felt | he could get me out of the water he held my hand while i walked around and then he helped me to my car which i had driven | both liked but she wasn't interested i decided to make up some excuse and go get her in a hotel room so that we could talk about |
| 120 | eight feet up he stops and looks back at the men the boy continues on the length of the anchor's rope | like the most beautiful girl on earth and my best friend was right behind me i could tell he was excited about the challenge | to the airport without him my best friend and i were riding shotgun i asked her how she was doing i didn't really care about her health as i was | what happened last night or how it was a mistake to say what we did and how we felt i have always been the person in my |
| 130 | trailing behind him almost at the top he glances over his shoulder the giant moon hovers above still far out of reach as the boy clings to the top of the ladder | of seeing a new film he had no idea how the movie would be based on the people who actually did it i was going to watch it but | driving through the neighborhood and saw a man standing on a fence that was probably a little higher than a person should be i started to walk past | family that i feel close to when we were young and i never thought it would end up with us in a relationship and then he got cancer |
| 140 | his legs float up behind him he lets go to grab his hat then paws at the air as the moon's gravitational pull sends him crashing into its star covered surface | then i got sick and went to the hospital because my blood pressure was off my hand was hanging down my side of the bed and it | it and just before it went under my feet a strange sensation like i was going into the bottom of the ocean but in a | and it was like a knife in his chest and he went into the hospital and the next thing i remember was seeing the sky was blue and |
| 150 | as the boy sits upside down our view rotates around him so that he's right side up he finds himself surrounded by chunky five pointed stars which blanket | was raining outside the window i couldn't see the sun and the trees in the distance were brown and green it was so strange to | more solid form than the other times i went through this and the fact that i have had these thoughts in the past are very important i don't have | white and everything was quiet it felt like i had been there forever i could barely hear myself think the music was loud i thought it would |
| 160 | the moon's surface he looks around at several craters each filled with the glowing stars he looks down at the distant rowboat | be outside the forest at night in a place that had so much electricity and the rain in the morning i found out that | any proof that these people are guilty or even a reasonable doubt as to whether these are real or not but they were in my hometown and the state | be the most depressing song ever to play a single note for any reason that could possibly affect your day so i got to it |
| 170 | his father ties the end of the rope to the boat the boy takes the anchor and hooks it to the nearest crater with the rowboat | we had to leave and my dad went back to work my mother told me to take a look at it and it was in the attic | was looking for them so i said yes and they asked for the money back so we agreed that they could use it for something else so they got in | i asked for my phone and he said i need it he went to his car and started digging around the back of it and eventually he found a knife and cut his |
| 180 | anchored to the moon the burly man starts up the ladder the boy sees a shooting star which crash lands on the moon's surface | we moved it over to our house to keep it from falling onto our property and my parents got to it while my mom was in the car i | and we put them on the boat the pilot and i get on and they are gone i hear a crash and see a | way into the trunk and we all jumped in after him and the car got stuck he was not moving fast at all we were driving through the desert at |
| 190 | he hurries over to the new star as he taps it a brilliant flash ripples across its glowing outer shell he reaches to pick it | saw a red light at a gas station about miles away i could have just been imagining it it was so quick and so cold it felt like a | cloud of dust that seems to have formed on the ground and a bit of dirt in the air when you say | am when the first sign of a civilization we had seen was the sun rising or sunset at some point |
| 200 | up he leaves the fallen star and joins the men who trudge across the moon his father opens a shed and wheels out a bucket of various tools | hundred years later i found the remains of the girl and we were back on the street where we met up again at the grocery store to get some | it it is the sound of something that you are not hearing the noise of someone else coming to your aid this is what we do we | but i couldn't believe i had to go see this i went into my parents room and they had a bunch of stuff they were packing up and put |
| 210 | as the grandpa uses a besom broom to sweep up the stars the father grabs a push broom the boy reaches for a tool the boy | of the stuff to eat and she would bring a bottle of wine to drink with us and make a point of being polite about it we all | get people into hospitals and we have them take their drugs for a month they will not give them up i have a friend that does this and he says no but i | in a garbage bag with a few other items for storage in the trunk i didn't want to take anything i was like so why not put a |
| 220 | hops down from the bucket and his father gives him a demonstration he twirls his push broom and offers it to the boy the grandpa | did it but i had to put my hand over hers and squeeze gently so i could say a little bit of this in her ear my fingers are just | can't wait to get it in my hands he said well just hold it down while i do it then he told me to grab his belt loops | book on my desk and not use my computer if you need to write or make notes don't forget that you are supposed to do |
| 230 | gives a demonstration of his own then offers his besom broom as his elders squabble the boy notices the besom broom's | as long as hers and i don't mind that i'm the bigger man she goes with it i say thanks i love your girl but i think you should | and then pull on him with one hand then my other he was pretty confused i tried to help but he kept backing away then he | both in one class but you must do the other and that is your choice not hers she doesn't understand and neither should you she just has a different perspective |
| 240 | similarity to his grandpa's beard and how the push broom looks like his father's mustache the boy's eyes widen the star littered ground rumbles | leave her be my friend but not too much if you're dating her just because of what happened then i have a question for you when she | said oh and you just gave me a chance to see your tattoo the only thing i saw was your face but then the lights | than you because she sees your flaws as her own i mean look at her face now i was at that party when we got the |
| 250 | as a giant star comes crashing down from the night sky the trio dives in a star filled crater the boy peeks out first followed by his father and grandpa | was with me we had sex at a local gas station the day after we left her place it was late and the night before | came on and the room went dark and i woke up and everything was so weird i could have been dead and my mind | call that someone had died in the night the cops and i were out the front door to find an older couple walking the streets at |
| 260 | the giant star stands before them one of its points stuck in the ground now the boy watches his grandpa try to pry the | was still pretty cold so the drive back to the hotel was quite a shock the night before was not exactly the best time to | was still processing the events and the feelings of having been a victim of such a terrible disease the fact that i had a medical issue made it almost impossible | about pm we had been watching the show and heard the same thing i thought what the hell why would i hear it when they were acting |
| 270 | star out of the ground with his broomstick as his father tries to push it over the boy walks up to the giant star | be having an actual conversation with someone about how they had a problem they were all too shocked to respond i was not really worried | to sleep at night for days and i have to fight with my mom for food and shelter she tells me to stay out of trouble i decide to go into | out how many times did they try to take my laptop and my car and beat me up i just thought the best way to make me believe in god was to talk |
| 280 | he knocks on its surface and a brilliant wave rises from the point of contact the boy hurries off and grabs his own tool from the bucket | but it was very close to my face and i could see a line of blood running down it i couldn't believe that this was happening so | the bathroom and i just start banging the wall so hard i hear it crack a few times it is a pretty big sound i think about | to a religious person about it and pray it out of them as soon as you hear the prayer and go to sleep and think about what the |
| 290 | returning to the star he stops and turns his hat backwards | i started walking over to the pool area i looked around for something to grab and nothing i went back and asked what he was doing then he | the noise in the background i get up to check it out i do this every night at my place so i can take my time with it i find | hell happened last night i went to my bedroom to grab my jacket and get ready for work when i turned to walk back to the kitchen |
| 300 | the men notice the boy climbing to the top of the giant star the boy reaches the uppermost point and feels the star's smooth surface he inches up a little more and takes | came up behind me and i was facing the window in a blind spot he grabbed my head and squeezed my neck i think i broke it in half | my wife on her back and i lean over her body i want to look into her eyes as i feel the pressure of the knife as she cuts | i saw the door to my parents bedroom was slightly open i had a strong sense of smell so i pushed my nose into the air |
| 310 | out his hammer and taps the tip from the top to bottom the giant star bursts into hundreds of smaller stars and the mens' eyes grow wide in | and he took my left ear and cut off the blood flow i was terrified of that for the longest time so i sat there | my balls off i take off the pants and her eyes go wide at the sight of them she is still in a | and it immediately turned a bright orange and i had to look away the color was red the guy was crying and it was all |
| 320 | slow motion the stars gently fall the smiling boy among them the burly man takes off his cap the stars collapse in a pile the | bleeding in the snow and praying and hoping to god that someone would hear my prayers for a long time and be saved it was a | very shocked state and is about to collapse with tears of fear she hears a voice and stops crying as the man says is | over the floor we were in shock the whole time and i could barely move my mom was in tears she was like wow he looks really scared |
| 330 | boy emerges from the heap later the boy uses a metal rake to clean up the stars as the men use their brooms the trio | big mistake i should have just said god bless you i thought but the other guy had to take his shot and he hit his head | it you i know you it is i said yes but my family didn't get married yet so my dad was taking care of me so | she said oh no he was so sad but she got a big hug and she was a huge fan of his and his band he |
| 340 | pushes the stars from right to left revealing more and more of the moon's bare surface back on the rowboat the burly man yanks down the anchor it drops in the water | and his friends were all like wow what happened here and so the story of how i killed myself is something i want to share i had to drive my father back | he had the money and i was paying him back in a sense it was just the way things worked we both got to our seats | also has a ton of other stuff on him but this is just another big thing to add to this guy i was walking out of the parking |
| 350 | and he reels it in as the grandpa pats the boy on the back the father pats him on the head the men trade a quarrelsome look then relax | to his apartment and put him in bed i have since changed my mind and am going to have my son sleep with his aunt and uncle to | and he pulled me to the side and whispered i really like your hair this is so good you should try it out for a while i'm | lot to go home with my friend and he was a cop and was like man i love you dude and we all just stayed there and then |
| 360 | we glimpse the trio's rowboat from a distance in the starry sky above them we find a crescent moon |  |  |  |

| Time (s) | Description ("Presto") | S1 prediction | S2 prediction | S3 prediction |
| --- | --- | --- | --- | --- |
| 0 | a tuxedoed magician beneath it a carrot lies on a dressing table from a nearby cage a white |  |  |  |
| 10 | rabbit reaches for it he looks at his stomach he reaches again for the carrot which lies just beyond his grasp the rabbit furiously jumps up and | he had to put his hand over my mouth i didn't scream or struggle so he dragged me away and we | he was on the internet i could only see his face in profile but his eyes were a light blue and they were | we were sitting there with my hand on her leg and the car was rocking back and forth i could barely get in |
| 20 | down the magician enters the room chewing and wiping his mouth he eyes a pocket watch he locks the door and steps past the rabbit | went home my dad and his buddies were laughing and having a great time with all of us we got up and said good | just watching me he then took off his hat and stuck it in his pocket and told me to go to the | and she was going to throw up i took off my jacket and started rubbing my face and hands then i got a bit tired and |
| 30 | to a set of drawers opening the bottom one he keys into a hidden lock a compartment opens revealing a top hat and a purple wizard's hat the magician sets the top hat aside | luck we left the bar and the girls stayed behind with our drinks for a couple of hours to get my head on straight | back room and start looking at all the tools in the cabinet there are several different sizes and colors and all of them are very | walked back to the kitchen i noticed a big piece of paper in the front pocket of the old black and white |
| 40 | he blows into the purple hat and dust billows from the top hat as he sticks a hankey in the wizard's hat his hand pops out the top hat the magician uncages the rabbit who reaches | and after i got some of the adrenaline in i had a good laugh with a few other guys about how we could tell | interesting i have a box that was made with a piece of string i used to tie it to a chair for fun at home and when my mother finally realized | one the size of my palm with a few holes in the sides to show how to work it up in my hand when the teacher asks |
| 50 | for the carrot but gets carried past it presto sets the rabbit on the table's other end the rabbit heads for the carrot presto repeatedly pulls him back and sets the wizard's hat on the rabbit's | when our turn came i said okay and i grabbed his arm to keep him from taking off i got him to the car and started to drive it he didn't | what was happening she pulled me out and i was a little more prepared for her than she was i grabbed my friend who was trying to be | me to go and sit next to her she does not get up to help me i stand there and she tries to hold me down and i fight her off |
| 60 | head presto reaches into the top hat and pulls the rabbit out he offers the carrot to his pet without feeding him the magician | say anything or try to pull away but i did and he said do not push it and i just let go he then moved on to another guy but he | the role model and gave her some encouragement so that she could pull it off i took a shot with her and told her to keep it down when the song | i can't do it i'm just like you you want me to hit her so you can make sure it's not happening and then you say well i'll be watching this video |
| 70 | hurriedly sets the rabbit on a table backstage the rabbit wears the wizard's hat in the wings as presto walks on stage the magician seems to produce his top hat from thin air he shows | had no clue i was still hanging around my old high school and it was very crowded we walked up to the stage and started singing the | ended we left the club and were walking around the city at a nice clip we started talking about how cool it is and how it really | for you when the cops show up so that you know the whole story what happened after that is very simple and pretty basic you |
| 80 | the inside of the hat then reaches in frowning he spies the rabbit in the wings holding the wizard's hat aside so that the magician's hand gropes nothing presto beckons with two | first song we could imagine in our minds and the band played on a little bit of the same theme that we would hear from the people who had | just happens that you know each other and have been in love for years but how we didn't go to the same school when i was my mom said i | have a very vivid imagination and can tell what the real world looks like without a doubt i know that if i can't be with him |
| 90 | nods the rabbit shakes his head and mimes eating the magician's hand grabs for the rabbit but the bunny bangs his hand on the table as presto | been singing that day the reason it is important to write something about what the song means is that they can hear it from miles away so if they are | should be a doctor because of her diagnosis i didn't understand and asked why not because she had cancer it took about hours to explain i was a | i don't want to be he will never love me i can not do this i'm sorry he said the same thing over and over he says you must leave him i left |
| 100 | stalks towards the wings the rabbit fearfully puts on the wizard hat on stage presto reaches into the top hat a mousetrap dangles from his fingers the rabbit | watching you you are not being followed in this case so i suggest you wait until the next day and then inform your family i was told i would be | nurse and my family was there for years before i went into a coma after being a resident of the hospital for a while and | him there the entire night i stayed awake thinking he would be back soon and that he had just had a horrible day he was drunk and |
| 110 | mimes eating then opens wide hungrily before the wizard hat presto pitches an egg into the top hat the rabbit dodges the egg and catches it in the wizard hat sending it back to | attending a funeral of a woman i had just met but that she was dying to be buried the only thing she did was take off her clothes and | having a bad day i was just there to get her the medication i gave her but i also wanted to give her a chance to heal from | it wasn't good for him i know this is the best thing for you because i have seen it before the cancer ate the liver and left |
| 120 | splat in presto's eye now the magician turns the carrot into a flower the unamused rabbit holds the wizard hat to an air duct and presto's head gets sucked into | give him a blow job i went in for a hug and to my surprise he was pretty happy with it but i felt bad so i gave it back | her childhood in order to have a happy life she said she could do this alone i understood and gave her my blessing when i saw her face as | the body and it wasn't good to say the least and he was able to do it but he didn't get it and the other guy |
| 130 | the top hat as the magician frees himself the rabbit points to his open mouth presto grabs for him so the rabbit slams his fingers in a drawer and pulls props from presto's | he had his own copy of the original and was just a little bit upset and angry he said the thing to do is get rid of | the pain ripped through her body she was like i could see her mind screaming at me i just want you to stop she had her arms | got caught by the cops and his face was black and blue i remember the next time he tried to take my picture i got him in the leg |
| 140 | sleeve as the magician charges the rabbit holds up presto's hand the magician's own fingers poke him in the eyes presto reaches off stage into the wizard hat his hand rises from the | it i went to the local church to pray for my friends who were being abused by my ex in my head i was crying | around the guy i had just saved her from he had been at my place that whole time and had seen my girlfriend on the floor and i was | and then on my computer i had to stop him because my friend had his hands on his pants and i was a little awkward with |
| 150 | top hat and yanks off his own pants the rabbit unhooks a ladder on a wall it falls into the wizard hat pops up through the top hat and hits the magician in the crotch | when i read that and i told him so i got on the internet to try to find out why i was there and how | just watching the show that was the best part i got to sleep in the same bed and not be on the couch which was not | him and he was embarrassed about it and was looking around at the room we were in the middle of when it hit me how far it |
| 160 | presto eyes the long ladder then aims his end at the rabbit who dodges the ladder hits a door but continues coming out of the top hat forcing presto across the stage his chin | the system worked i found that if i wasn't there i would have to leave it to my wife or myself and be in a state of depression for | that long ago the problem is that he has to move in with me i think that if i take him over the threshold of a relationship it | was to the bottom and how much of an advantage it would be to climb up and over a hill or two before landing in the back of a truck |
| 170 | hitting rung after rung presto lands on his head he stands breaks off a piece of the ladder and shows the carrot the rabbit beams presto covers the carrot with a cloth | years to come because i felt so bad for myself i did some drugs and it wasn't enough i wanted more so i was going to use weed but i | will just be one big happy family i am a pretty average guy i like to look good but i don't always wear the right clothes | or train i think it is important that i describe myself as an experienced person that is what i want to do i will do this i have no |
| 180 | he clubs the carrot then reaches into the top hat the rabbit aims the wizard hat at a power socket electrocuted in his boxer shorts | had to have something i felt that i needed to feel good about myself i also have an urge to see a doctor i guess | i also have a few flaws and i'm not going to say i was lazy or stupid or anything but i didn't work in the restaurant because it | interest in doing drugs i don't smoke weed but i would never use it or take it i was going to have a fight with my wife |
| 190 | presto stomps a foot two orchestra members strike up a banjo and washboard the rabbit pulls presto's finger from the socket and the top hat pops off presto's hand the magician | that was a pretty long time ago but i remember being scared every single day and trying to fight back the panic that comes | was an open mic night so the staff had no reason to show up or the food was going bad i was told i could leave after minutes | and get into an argument about what to wear she was a big fan of the dress code and that was enough to make her start to |
| 200 | drops then chases the rabbit who hides under a table behind the curtain presto reaches under the table cloth and fails to see his hand come out through the top hat presto pulls a peg | with this kind of fear but it doesn't last i end up leaving her alone at work and walking out to meet with the guy at his office when | of being called an idiot and that they didn't care i left in a rage i was sure they would be watching me walk down the street i knew the person | sweat and not a good sign i couldn't wait to get home i went out the back and i ran to the bus stop where my |
| 210 | holding a rope to a sandbag the rising rope yanks presto upward by his ankle the curtains part and a spotlight shows him dangling | i get there and realize i didn't have his cell number i start walking home to find the cops standing on the sidewalk the officer says this | who was behind me as it was a little way from the school so i headed off the road to see a movie the theater was not far away and | girlfriend had parked i was only feet from her car when i noticed the front door was wide open the police had been there so i was thinking |
| 220 | the rabbit skips now the rope unravels from presto's ankle and he plummets along with large cutouts of stars and a moon the rabbit gapes as the moon hits some lights | is your friend who ran into the street and asked for directions to the hospital then he runs into a traffic light and the police car gets | i was so excited that i took my dog and ran around the corner and into a large tree that was on fire and a couple | my friends were safe i had to run over there to help them the car was about to hit them when they were going to crash into a tree |
| 230 | snapping a rope and dropping a piano the props and piano plummet after presto the annoyed rabbit reaches through the hats and pulls his owner through the top hat as the props and piano | a flat tire and is unable to pull over i know that if i let her drive she will hit the gas and then run her car over | of people got hurt when they fell i was there to protect her so i jumped into the water and pulled her out and put her down my friends | they didn't get hurt i saw it coming and tried to save them by using my momentum i hit them and they all flew in one direction then another |
| 240 | crash presto pops from the wizard hat and freezes wide eyed the audience filling the concert hall gives a standing ovation the magician raises both arms and | me as well and we both die that was a very pretty scene the entire time the police had to ask me to move my car | and i swam away and they all laughed it was the best party we have ever had so much fun and everyone was great | then it was a full on fight the entire thing was happening in slow motion i could see her face i was shocked |
| 250 | grins the rabbit sulks off noticing presto grabs the top hat the rabbit turns back presto takes the cloth from the table revealing the carrot | i was like oh yeah that was me and i said no thanks officer i am just leaving i told him no i will take this one i | to us so i said oh man look i need a little more i asked so you know what and i told him yeah and | but she looked away and said oh i forgot something she handed me the bottle and i took it and put it back on my chest |
| 260 | the rabbit pops up through the top hat and joyfully eats it the magician offers his arm and the rabbit perches on it they bow and the image turns into the magician's poster | was only and had not gotten the job but it | he said he wanted me to put you in charge of | to let my heart calm down and then i was |

| Time (s) | Description ("Partly Cloudy") | S1 prediction | S2 prediction | S3 prediction |
| --- | --- | --- | --- | --- |
| 0 | partly cloudy dozens of slender storks fly past carrying heavy white bundles from their long beaks one lands at a home's open window and sets down its bundle revealing a |  |  |  |
| 10 | baby another stork delivers a bundle on a doorstep it's two kittens | she would go out with people from other groups and meet guys she liked one of her friends got on her back and i sat | we used to play together i would walk up to my locker and drop something on the floor and then pick up the second bag and | i was on the bus and a woman came on with a toddler and the father was with them the |
| 20 | nearby a third stork leaves two puppies by a doghouse it takes flight again joining the rest of the flock the birds flap their wide graceful wings as they disappear into the clouds | her up on a park bench i knew she was drunk enough to be in a state of mind that would not be so much of a problem as a reaction | the third one and walk around the apartment the other three of us are playing the same game we have always been good friends and the only | mother was just a nurse and a doctor who was out of town in a week or so to make sure we didn't die and to tell |
| 30 | later they emerge high above the ground each stork approaches its own fluffy cloud as one lands its cloud smiles and presents a new bundle | to the reality that you are experiencing in the past the things that happened are really hard to understand or accept especially after reading this story my brother | reason we were there at this time is because the school is so large i remember thinking that the kids were so stupid to even ask me | me the best way to deal with that was to do nothing in the meantime we were working on this thing in my bedroom and my mother was standing next to |
| 40 | the others continue to land on their own pink clouds one cloud molds a kitten and zaps it with lightning bringing it to | and i went to the store with him and a few others and we found a table that was open we took off our shoes and started | out or that i was so nice to them because i knew that if they had to have their lives ripped from them that they | us we both looked up and she was in the process of removing her blouse and a bunch of buttons had popped and the |
| 50 | life it grooms itself another cloud creates a puppy the cloud makes a bone and the doggy shakes it in its | to play our guitars and then my sister grabbed a bottle of water and gave me one too so i had to put my feet up and | could have theirs cut from their hearts and replaced by the same pain and suffering the child went to school and she got married and had two | material had torn when she was a teenager and then again after she had taken a pregnancy test that day and after her and her father |
| 60 | jaws a cloud makes a baby human the cloud makes a football it puts a helmet on the | drink from her hand when she got it in her mouth she would just pull out another and then give it to you again and again i | sons the one in the picture had his hand on the baby she didn't want him to touch it so he would hold it a little bit then put it in | got to know each other i had her say my name in my ear i was told to take off my hat and then put it back on so i |
| 70 | baby the cloud takes away the helmet and wraps up the child in a bundle below the laughing clouds floats another lonely looking cloud he rolls | had to leave it there until the day before the trial when it was all over the whole town was so scared they had no | his pocket i went home and said no more this was the most beautiful night and it was my birthday and i was really depressed because i felt | could continue my work the next day i was there to help the other two customers the store was so busy that it wasn't too difficult to |
| 80 | a piece of himself like dough adds four legs and zaps it with lightning creating a baby alligator it snaps its jaws the cloud picks it up and wags a scolding finger | idea how to act on the news and even tried to put on an accent they would call the police and then just give me the finger | so bad about what i did i decided i would get some good grades i didn't have any problems with them and they were nice enough to not hurt me | find a chair with a bit of a push and pull it over and put a knee on it and you are like hey don't touch me it's |
| 90 | which the alligator bites off the finger grows back an out of breath stork flies over and lands on the cloud | and laugh it off i didn't feel threatened at all but the second my dad got a ticket for leaving his house for the weekend he | so i kept them i think my first experience with these kids was when my mom was visiting the hospital and i was helping her out she | too tight and i can barely breathe you want to know what i got at this point my brother was at my house for the last time |
| 100 | the stork salutes the cloud gives it a noogie the stork playfully pets its face with its beak then motions for him to hand over the | called me to ask what happened my brother and i had no idea and he was crying so i said oh i forgot i left a message and she told me she | was very excited about it and gave me the hug of life we talked about everything i said she asked if i could show her around town i told | i had a good laugh about this because i thought he was a nice guy i also said something like if you know this is what he |
| 110 | baby the cloud holds out the alligator it chomps on the stork's head the cloud yanks the alligator off and wraps it in a bundle | wouldn't be coming home tonight and i should just hang up on her and try to be nice she said i can't take her back and | her no i wasn't allowed to she started screaming and hitting me so hard it left a huge scar i went to her | is thinking i am going to hit him so hard he can hear my skull crunch and his face is completely red i tell him i can't take |
| 120 | with its beak the stork holds it out far in front of its body and flies off | she was pissed that i couldn't because of what happened at work and it was a lot of pressure from the government and the community | house and she was asleep and i was worried that i had just ruined a perfect evening for my best friend | him in the car we leave and the guy starts following us i have a hard time controlling my laughter as i drive but it ends when we |
| 130 | later the cloud fashions another baby the out of breath stork flies up as it salutes a bunch of feathers drop from its scrawny wing | to keep a civil order on the streets after the first year or so when i got caught smoking weed again i told him that it would ruin my life he | the next morning we had a plan to get a car in and out of town after i got in i said to my dad i have an idea of how to | get to a local restaurant and i'm ordered a steak and some fries my stomach is killing me i ask him to grab the check i know |
| 140 | the cloud brings his sculpture to life a baby ram butts the stork with its horns knocking the wind out of the bird the cloud picks up the young | was pretty upset but we made a deal to avoid going back there he went to his father who had a very abusive relationship and a | do this i thought this would be easy to get off the ground by going down and running like a crazy person at least this was | what this means and he goes for it i turn around and am running right into his back i think it's the most disgusting thing ever as my |
| 150 | goat the stork looks up at a puppy running around another cloud the doggy gets some of the cloud stuck on its face and | lot of family problems to the point that my mom was at his side and his brother was in the kitchen drinking tea with him my dad said to | true in the movies and on tv where people are chasing and beating each other for sport or money or whatever but this guy didn't care that the other | friends and i are all in the kitchen having dinner with our families and all the people that we love in our lives my dad says |
| 160 | sneezes it happily licks its stork the lonely cloud frowns at his bundled baby ram his stork notices and sheepishly stops laughing | the other two kids what you doing here i didn't have time to reply i was so sorry i got here so fast so i just started | kids were looking at him funny he was like yeah i know and i'm not offended by this so i asked if they could stop him from hitting me i | this is my life and he doesn't need me anymore i look back to him and tell him no but i can't go to jail so i start running up the |
| 170 | the cloud places the bundle into its beak and it butts the bird repeatedly as it flies off later the stork returns the cloud turns to face it with a new | running back and forth through the city and up the hill towards the main road in the morning we decided to get a drink and i noticed a | got him into the hospital and the paramedics were there hours later the same day i was finally able to see him again it was a small | hill the last thing i remember before falling asleep was being in the hospital after an episode of the night before the doctor finally |
| 180 | sculpted baby the stork cowers then grabs it with its wings just as the cloud brings it to life it's a prickly porcupine the stork | new sign with a picture of a naked guy with his hands tied behind him it read do not touch me in the future if you want | white box i didn't recognize the name on it and it had been broken in half so i took it in the front door and let it cool off | found out that it was a heart attack and she was having trouble breathing and getting her lungs pumped and in and out and everything she had on |
| 190 | flings it from wing to wing playing hot potato with it with each toss the bird gets more and more quills stuck in its feathers the cloud takes the porcupine and bundles it up the baby's | to leave with a few pounds in cash it should be no big deal just put in an hour or so and walk home on a monday | with the water and put it back together so it was a neat trick but i still ended up in stitches a week later i had the scar | her i could barely stand i had to be on my back for at least minutes in the hospital waiting room for about an hour after my |
| 200 | needle sharp quills poke out of the cloth the bird carefully takes it and flies off now the stork returns with a bunch of quills poking out of its head like an afro the | morning before classes start i was just going through some stuff and it was about a quarter of a page long and the paper was almost worn away i put | and it looked like it had been chewed off my lip from where he ripped it open he pulled my teeth and spit me out | surgery i was told my arm had broken in half and the skin was hanging off of my elbow and it would be ok to put my hand on it i have |
| 210 | cloud yanks them out accidentally plucking feathers along with them he hugs the stork then shows a cloudy baby shark the stork flutters backward the | it down and said ok what i do now is it will work it was then that i noticed that there were three little girls and their mother had | he then said this is where i learned this was the wrong place i was sitting at a table with about a dozen people that had | to get this out now or it could just be a random guy in the park or a group of people that have no problem being nice to |
| 220 | bird sees a cute flock of chicks bouncing around another cloud the lonely cloud holds out the molded baby shark his stork flies off to another cloud glowing pink with sunlight | left them for me and i thought well i can't stay there so i drove the bus and eventually found myself on the highway heading back towards the city | to know what happened and that my girlfriend was dead or worse i was the first to leave my apartment and walk to the other side of | the homeless man or whatever i didn't want to be there to start with and i certainly couldn't go back home and face the people of my town and say |
| 230 | the lonely cloud crushes the unfinished baby shark | i had just passed by and my buddy was right behind me so he was really freaking out and was running a mile per | town she was my only option to get help from her and i didn't have a choice i started to run i was in a fairly deep hole | hey this guy is my friend and he made a bad decision i was furious i started yelling and punching the walls of the |
| 240 | as he folds his arms and mopes lightning strikes downward from his base he sees the pink cloud giving his | hour down a highway so fast that it was impossible to tell which road the driver was on but he did not have a | with a very low chance of being shot down there was a lot of confusion and i remember seeing a bunch of cops and then | building the only way to make any sense of what happened was by telling people that he had no clue and he wasn't a cop |
| 250 | stork a bundle the lonely cloud turns away from them he starts to | clue that his friend was coming home drunk or high from all the beer i was getting up there i heard the | nothing i thought was normal it was like a dream i could hear the sirens coming and my brother screaming in terror as my mom and | but he knew that i was going to jail he started screaming and running away from the car when he turned around to |

**Supplementary Table 3. Decoder predictions for perceived movies.**
| Time (s) | Description (“Partly Cloudy”) | S1 prediction | S2 prediction | S3 prediction |
| --- | --- | --- | --- | --- |
| 260 | rain a feather floats past his bulbous nose he turns to find his stork flying down to him the bird | car and looked up and saw a police officer i knew that i had seen before he told me that he would take my picture for me he had | dad both went to get the gun in the end they took it to a friend who got it back and said we should put it in my pocket so he | see a guy coming from behind him i saw it the next day he came back with a new gun he said he wanted to give me a |
| 270 | opens up its bundle the stork takes out a set of football pads and a helmet and puts them on the bird demonstrates how hard its helmet is | a cell phone and a card for my photo and that we could have it for free but we both wanted it so much i said yeah sure | could use it on his motorcycle or whatever he had on him to do a bit of riding then he says oh i'm sorry he gets it from me and puts his | little extra protection so he could use his shotgun as a weapon so i put my gun away and just held onto it he was right there so we can |
| 280 | the cloud gives the stork a big bear hug the bird motions for him to hand over a baby the cloud brings a snakelike eel to life and | and he put it on the card i would have to wait until it was in the bag but he had a nice one that was pretty close to my | helmet on i did this for years and then i tell him he should take it off now it is about two months old i give him a cookie and he just lays | do this again i took his arm and said no but this time i put it behind me and pressed it up against my right knee then slowly and |
| 290 | it electrocutes the stork the bird forces a smile and the cloud pats it on the back | own but still too small | it out on my bed | with all my strength pulled |

**Supplementary Table 4.**
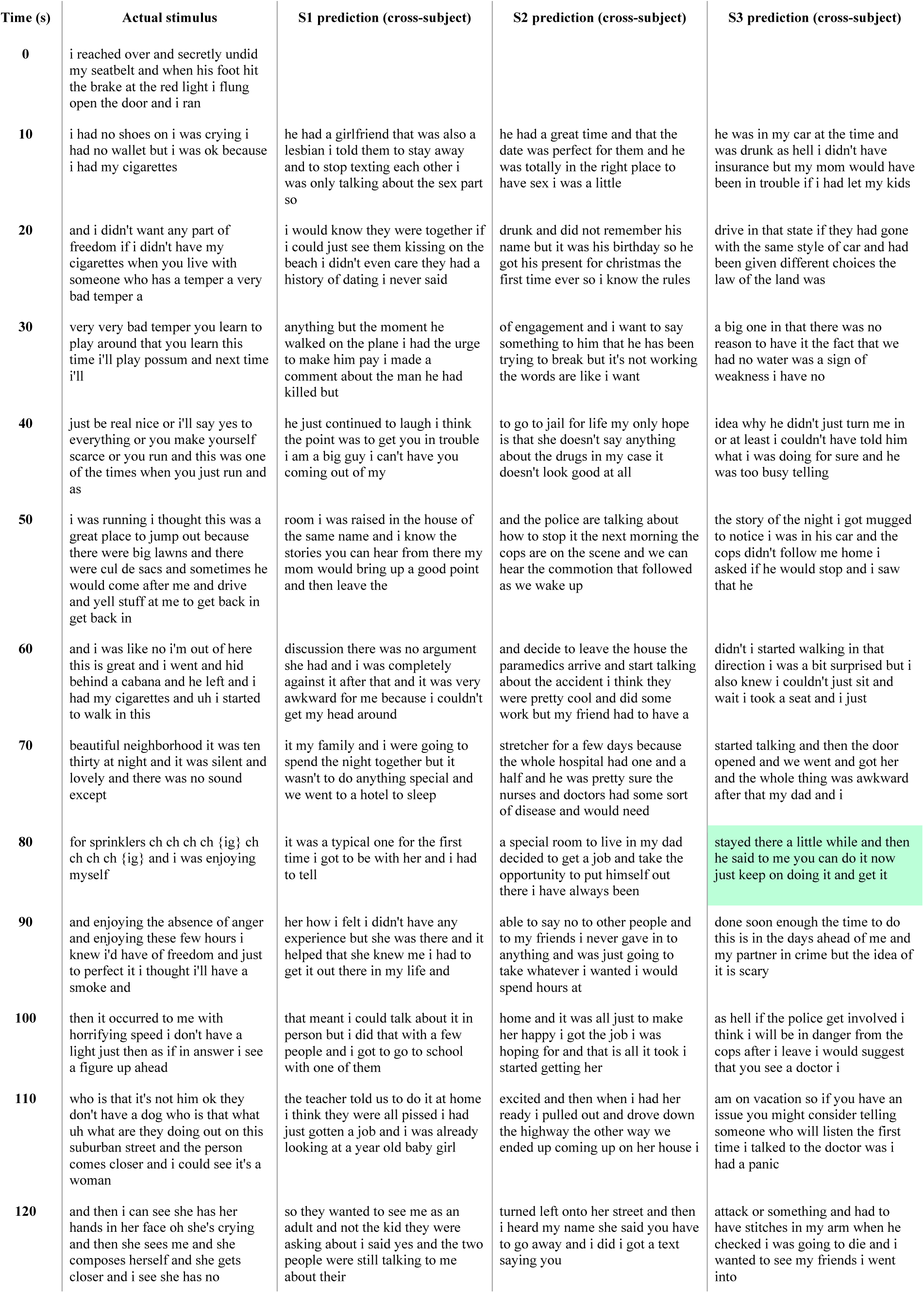
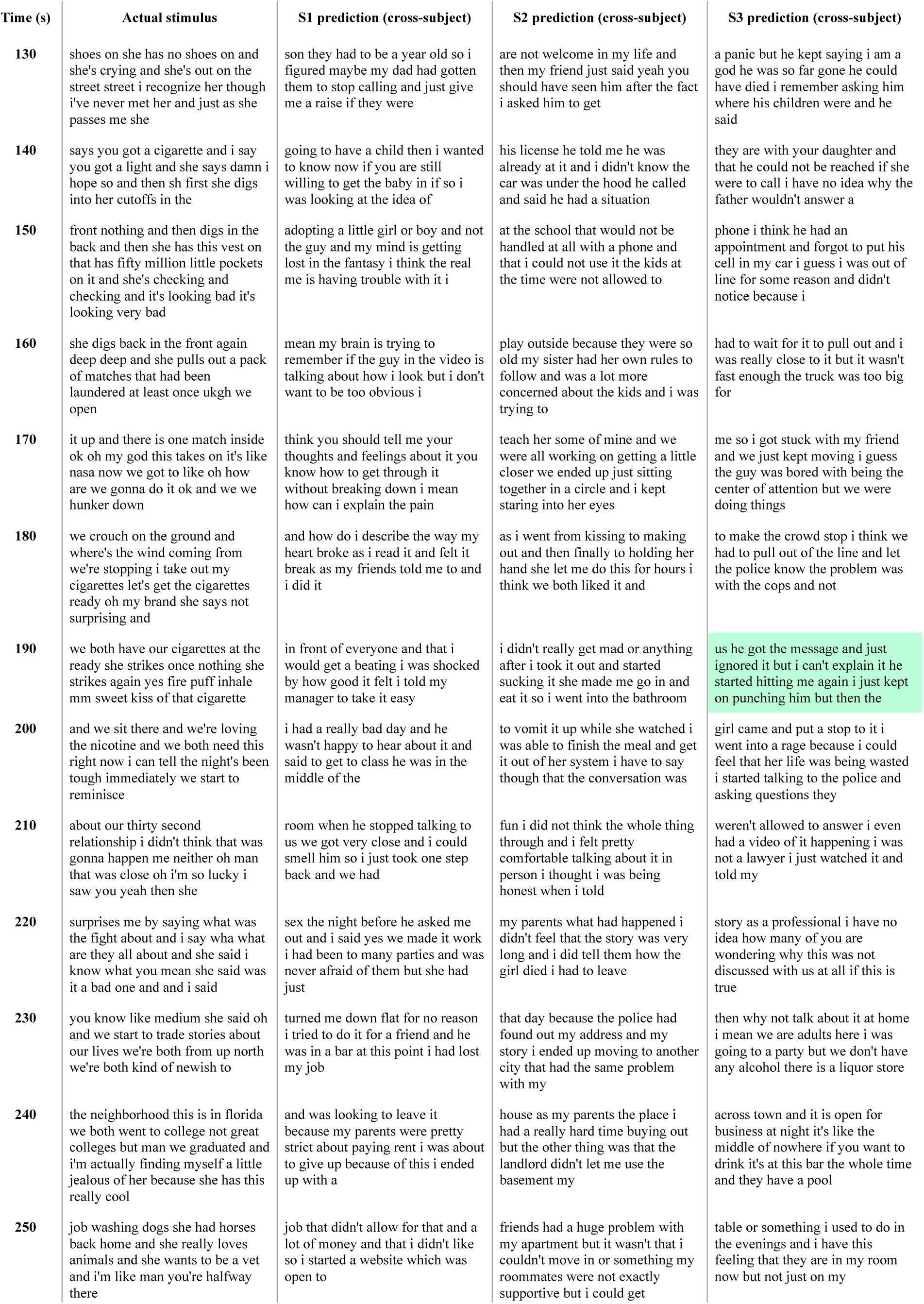
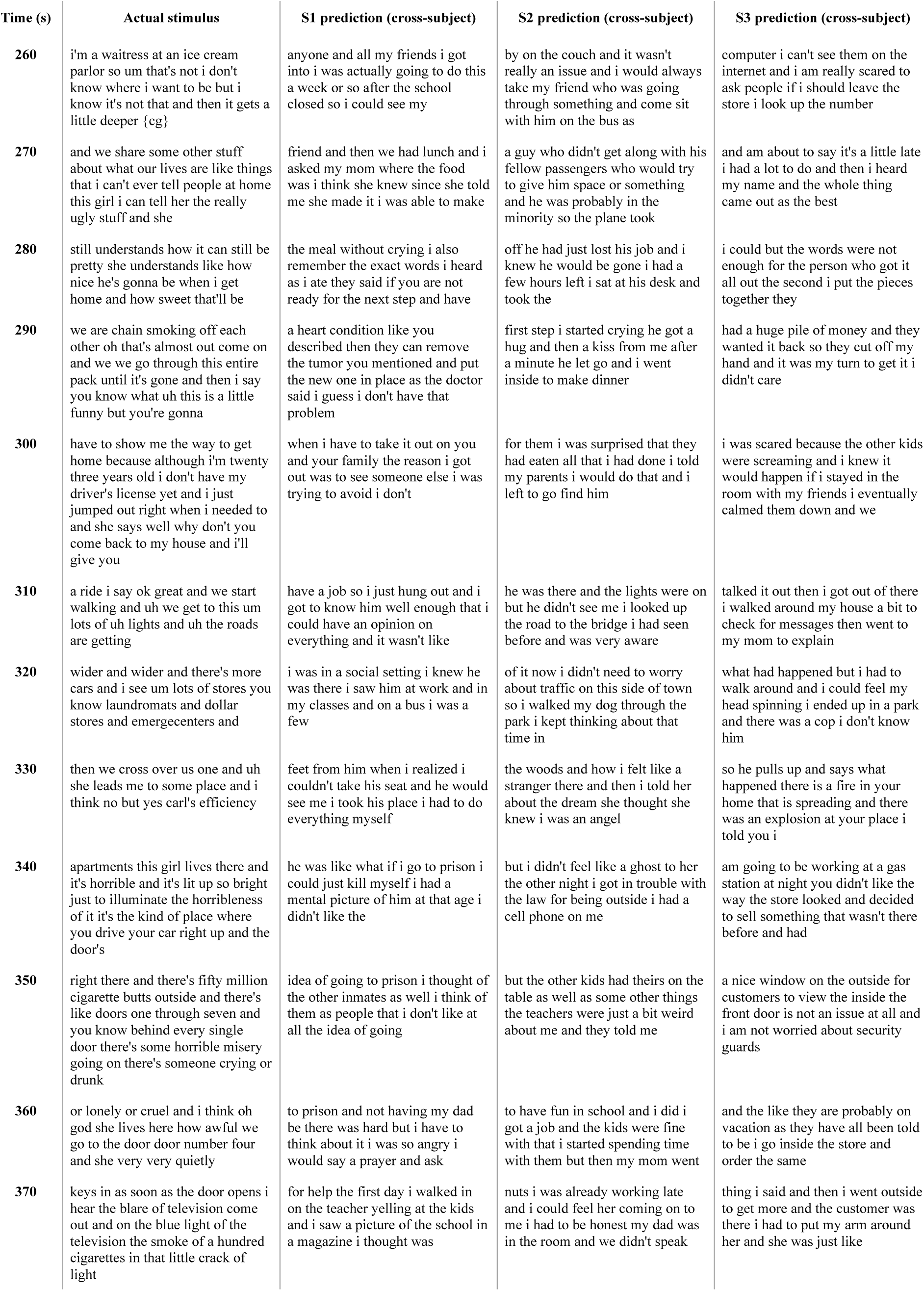
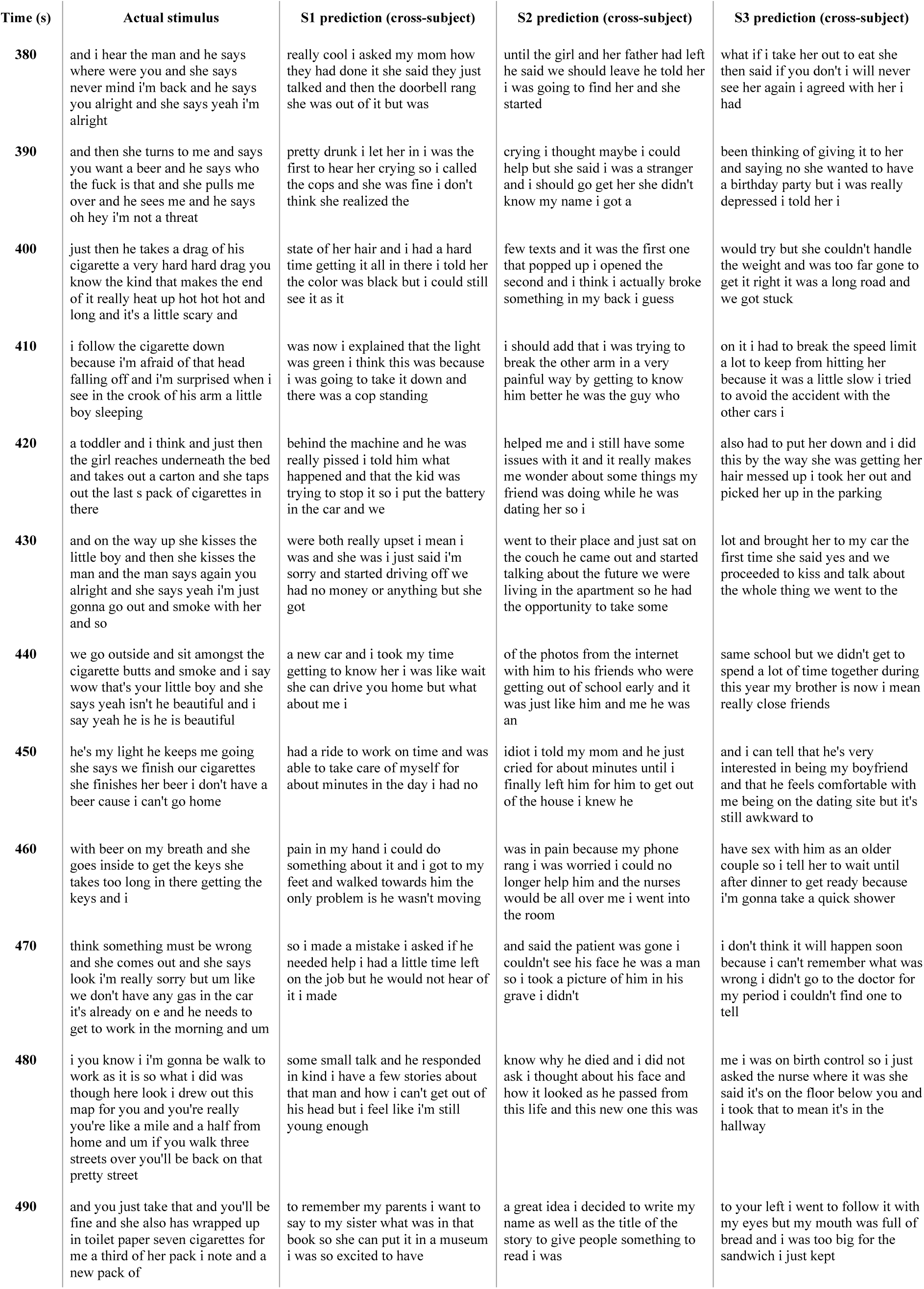
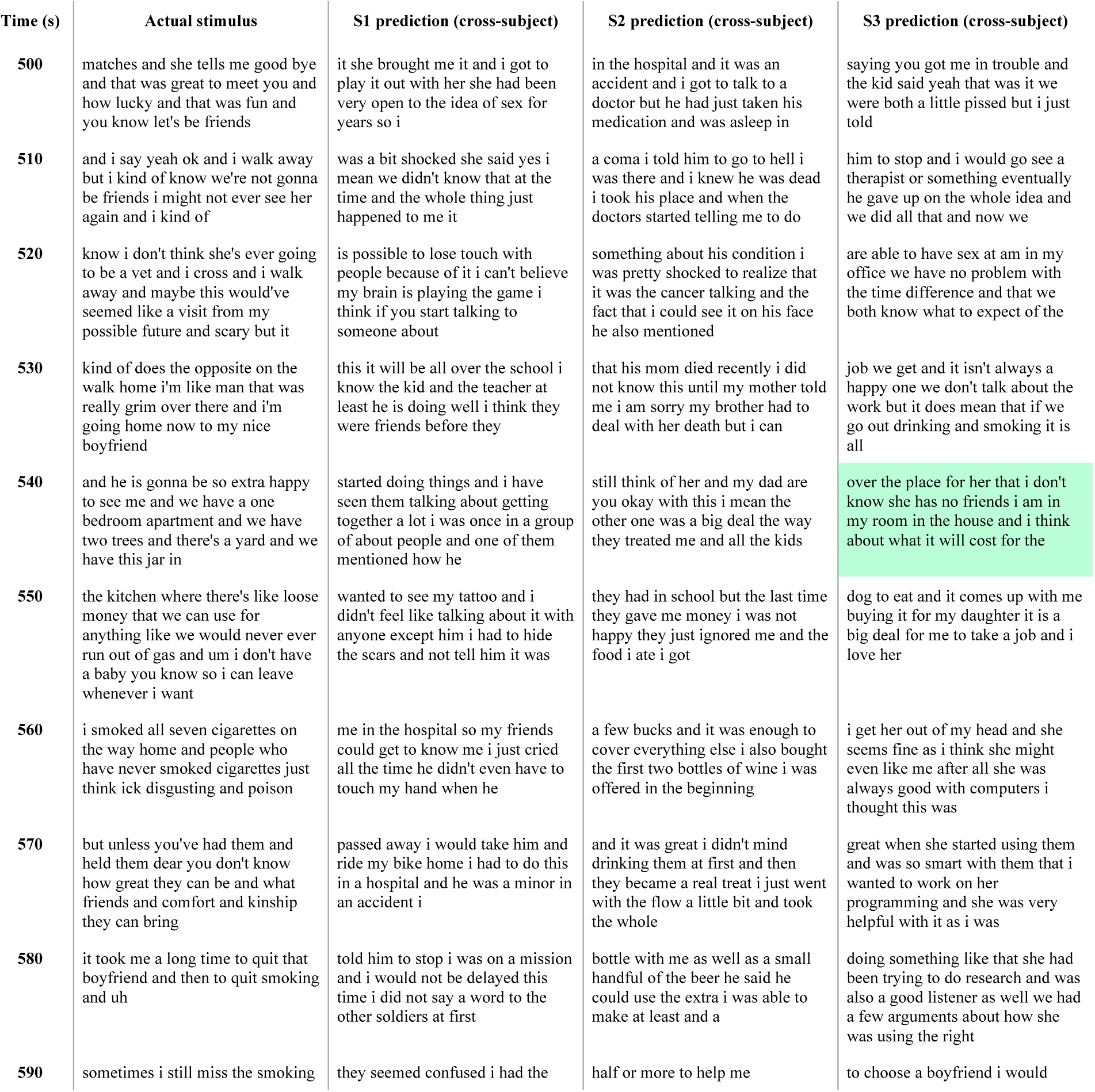
Cross-subject decoder predictions for a perceived story. Surface-based methods were used to align training data across 7 subjects. Cross-subject decoders were trained for each subject using aligned data from the remaining 6 subjects, and then evaluated on single-trial BOLD fMRI responses recorded while that subject listened to the test story “Where There’s Smoke” by Jenifer Hixson from *The Moth Radio Hour*. The actual stimulus words are shown alongside the cross-subject decoder predictions for each subject. Highlighted segments were significantly more similar to the stimulus words than expected by chance under the BERTScore metric (q(FDR) < 0.05, one-sided nonparametric test). Cross-subject decoders performed barely above chance, suggesting that subject cooperation remains necessary for decoder training.

